# *Gastruloids* develop the three body axes in the absence of extraembryonic tissues and spatially localised signalling

**DOI:** 10.1101/104539

**Authors:** D.A. Turner, L. Alonso-Crisostomo, M. Girgin, P. Baillie-Johnson, C. R. Glodowski, P. C. Hayward, J. Collignon, C. Gustavsen, P. Serup, B. Steventon, M. Lutolf, A. Martinez Arias

## Abstract

Establishment of the three body axes is a critical step during animal development. In mammals, genetic studies have shown that a combination of precisely deployed signals from extraembryonic tissues position the anteroposterior axis (AP) within the embryo and lead to the emergence of the dorsoventral (DV) and left-right (LR) axes. We have used *Gastruloids*, embryonic organoids, as a model system to understand this process and find that they are able to develop AP, DV and LR axes as well as to undergo axial elongation in a manner that mirror embryos. The *Gastruloids* can be grown for 160 hours and form derivatives from ectoderm, mesoderm and endoderm. We focus on the AP axis and show that in the *Gastruloids* this axis is registered in the expression of T/Bra at one pole that corresponds to the tip of the elongation. We find that localisation of T/Bra expression depends on the combined activities of Wnt/*β*-Catenin and Nodal/Smad2,3 signalling, and that BMP signalling is dispensable for this process. Furthermore, AP axis specification occurs in the absence of both extraembryonic tissues and of localised sources of signalling. Our experiments show that Nodal, together with Wnt/*β*-Catenin signalling, is essential for the expression of T/Bra but that Wnt signalling has a separable activity in the elongation of the axis. The results lead us to suggest that, in the embryo, the role of the extraembryonic tissues might not be to induce the axes but to bias an intrinsic ability of the embryo to break its initial symmetry and organise its axes.

**One sentence summary:** Culture of aggregates of defined number of Embryonic Stem cells leads to self-organised embryo-like structures which, in the absence of localised signalling from extra embryonic tissues and under the autonomous influence of Wnt and Nodal signalling, develop the three main axes of the body.

## 1 Introduction

The establishment of the anteroposterior (AP) and dorsoventral (DV) axes during the early stages of animal development is a fundamental patterning event that guides the spatial organisation of tissues and organs. Although this process differs from one organism to another, in all cases it involves a break in an initial molecular or cellular symmetry resulting in the precise positioning of signalling centres that will drive subsequent patterning events. Dipteran and avian embryos provide extreme examples of the strategies associated with these processes. For example, in *Drosophila*, the symmetry is broken before fertilisation within a single cell, the oocyte, which acquires information for both the AP and DV axes. This occurs through interactions with surrounding support cells that control processes of RNA and protein localisation which then serve as references for the rapid patterning of the embryo as the zygote turns into a multicellular system (2, 3). On the other hand, in chickens the processes take place in the developing embryo, within a homogeneous multicellular system that lacks external references (4, 5). In mammalian embryos, the axes are also established within a homogeneous cellular system, the epiblast, but in this case they are under the influence of an initial symmetry breaking event that takes place within the extraembryonic tissues which is then transferred to the developing embryo (5–8).

Efforts to understand the molecular mechanisms that pattern early embryos have relied on genetic approaches such as perturbation of the process by genetic mutations and a correlation between specific processes and molecular events as highlighted by the genes (9, 10). Although successful, these approaches have limitations as they often conflate correlation and causation, and importantly cannot probe the role of mechanical forces that have been shown to play a role in the early events (11, 12). This suggests a need for a complementary experimental system in which, for example, rather than removing components we attempt to build tissues and organs from cells and learn what the minimal conditions are that allow this (13). We have recently established a non-adherent culture system for mouse Embryonic Stem Cells (ESCs) in which small aggregates of defined numbers of cells undergo symmetry breaking, polarisation of gene expression and axial development in a reproducible manner that mirrors events in embryos (14–16). We call these polarised aggregates *Gastruloids* and have suggested that they provide a versatile and useful system to analyse the mechanisms that mediate cell fate assignments and pattern formation in mammalian embryos (16, 17).

In this study we show that *Gastruloids* develop the three main axes of a mouse embryo (AP, DV and left-right (LR)) in the absence of extraembryonic tissues and investigate the underlying molecular causes for this event. Unlike the embryo, this process does not require BMP signalling but relies on interactions between Nodal and Wnt signalling that are recorded in the expression of the transcription factor T/Brachyury (T/Bra). Furthermore we show that localisation of Nodal, which is widely held as essential for the establishment of the AP axis, is not required for the polarisation of T/Bra expression. Our results provide novel insights into the patterning of the embryo and suggest that a similar spontaneous symmetry breaking event may occur in the embryo where biases from the extraembryonic tissues, ensure its reproducible location at the site of gastrulation.

## 2 Results

### 2.1 Axial organisation in Gastruloids

Our previous studies using *Gastruloids* revealed them to have an AP axis with the expression of T/Bra located towards one end that will lead an elongation process (14, 16, 18). To follow on from these observations and to determine whether other markers of the embryonic axis are present, we cultured *Gastruloids* for 120h, probed for a range of axis-specific markers by immunofluorescence, and mapped the expression domain of reporters for the two major signalling pathways involved in axial organisation in the embryo: Wnt/*β*-catenin (Tcf/LEF) and Nodal (Smad2,3) signalling (Fig. 1 and g. S1 and Materials and Methods). At 120h After Aggregation (AA) *Gastruloids*, which have been exposed to the Wnt signalling agonist CHI99201 (Chi) (19) between 48 and 72h AA, are clearly polarised, with localised expression of T/Bra (Fig. 1A) and Cdx2 (fig. S1A) at one end of the protruding tip. They also exhibit a shallow gradient of Wnt signalling away from the T/Bra expressing region (Fig. 1C top and insert; Fig. S1C, top). The positioning of T/Bra expression suggests that this region is similar to the tail bud of an embryo supporting our previous observations that *Gastruloids* have AP axial organisation (20–22) where T/Bra and Cdx2 define the posterior domain. Correlating the expression of Sox2, a Sox1::GFP reporter (23) and a Nodal::YFP transcriptional reporter (24) (Fig. 1A, Fig. 1B, fig. S1A), we observe an additional, perpendicular axis where high levels of expression of the neural development markers Sox1 and Sox2 extend away from the T/Bra-expressing tip on one side of the *Gastruloid* and a tight clustering of Nodal::YFP expression directly opposite (Fig. 1A, Fig. 1B; fig. S1A). This second, orthogonal axis has a strong similarity with the DV axis of the early embryo, and suggests that *Gastruloids* are capable of generating a Node-like structure (Fig. 1C (bottom); fig. S1B, fig. S1C), a conjecture supported by two observations. Firstly, the ventral expression of Nodal::YFP exhibits a weak but noticeable asymmetric expression similar to that of the Smad2,3 reporter (Fig. 1C, middle and lower; Fig. 1D, top). Secondly, the Smad2,3 activity around the putative node-like region evolves from a symmetric to a bilaterally asymmetry emerging over time (Fig. 1C; top, middle and insert and Fig. 1D). Shaking of the *Gastruloids* from 120h AA allows the cultures to proceed until 160h AA and reveal a more complex organisation in coherent structures over 1mm in length (Fig. 2). The AP and DV organisations are maintained and the Sox1 expressing tissue exhibits a bent morpholoy reminiscent of the organisation of the developing spinal cord (Fig. 2B). Inside, we observe a long tubular epithelium (Fig. 2B, C); these cells express low levels of Sox2 and high levels of Sox17 and therefore are likely to correspond to endodermal derivatives.

**Figure 1:**
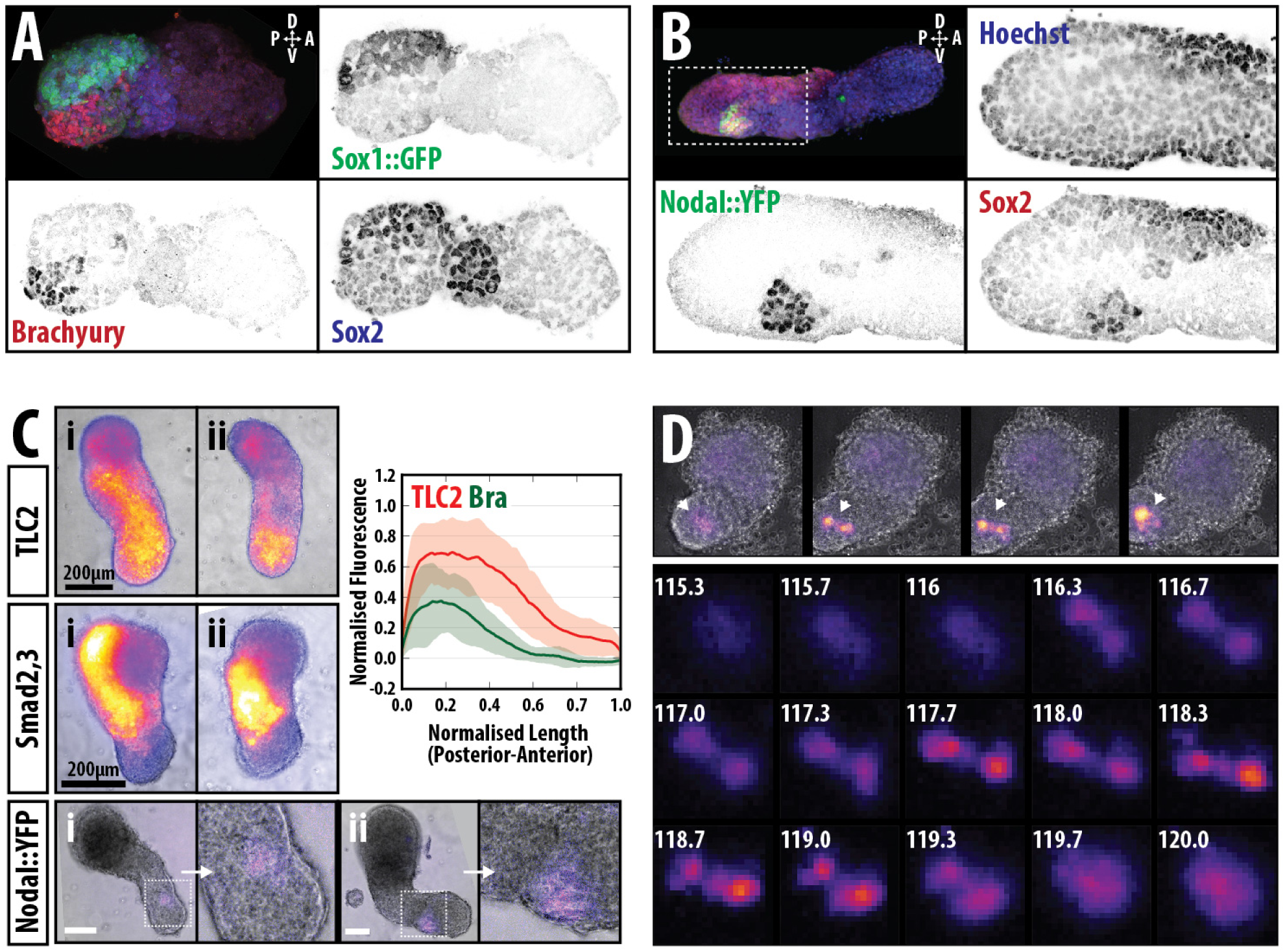
The anteroposterior, dorsoventral and left-right axes are clearly defined in *Gastruloids* and develop a Node-like region. Sox1::GFP (A) and Nodal::YFP reporter (B) *Gastruloids* pulsed with Chi (48-72hAA) and stained with Hoechst and anti-GFP with either (A) T/Bra (red) and Sox2 (Blue) or (B) Sox2 (red) at 120h AA. Hoechst is not shown in (A). Nodal is clearly expressed opposite an extending plate of Sox2 expression and T/Bra is observed at the most posterior region of the *Gastruloid*. In both examples, 3D projections are displayed. Taken in combination with supplemental figure S1, the expression of different genes and reporters can be used to map axes of the embryo onto the *Gastruloid*. (C) *Gastruloids* formed from Wnt/*β*-Catenin (TLC2), Smad2/3 and Nodal::YFP reporters lines following a 48-72h Chi pulse are shown. Insert shows the quantification of reporter expression for the TLC2 (red) and Bra::GFP (green) *Gastruloids* in a posterior to anterior direction. Stimulation results in activation of the TLC2 reporter with highest expression at the posterior pole. Interestingly, lateral expression of the Smad2/3 reporter is observed in a large proportion of *Gastruloids*, indicating the initiation of a LR axis. Nodal expression by 120h is confined to the posterior region indicating a prospective Node-like region. (D) A small number of Smad2/3 reporter *Gastruloids* at 120h show expression in two regions at the elongated tip at a low frequency. Over time one side is down-regulated similar to the expression of Nodal in the perinodal crown cells of the embryo (see Fig. 1C&E in ref (1).

**Figure 2:**
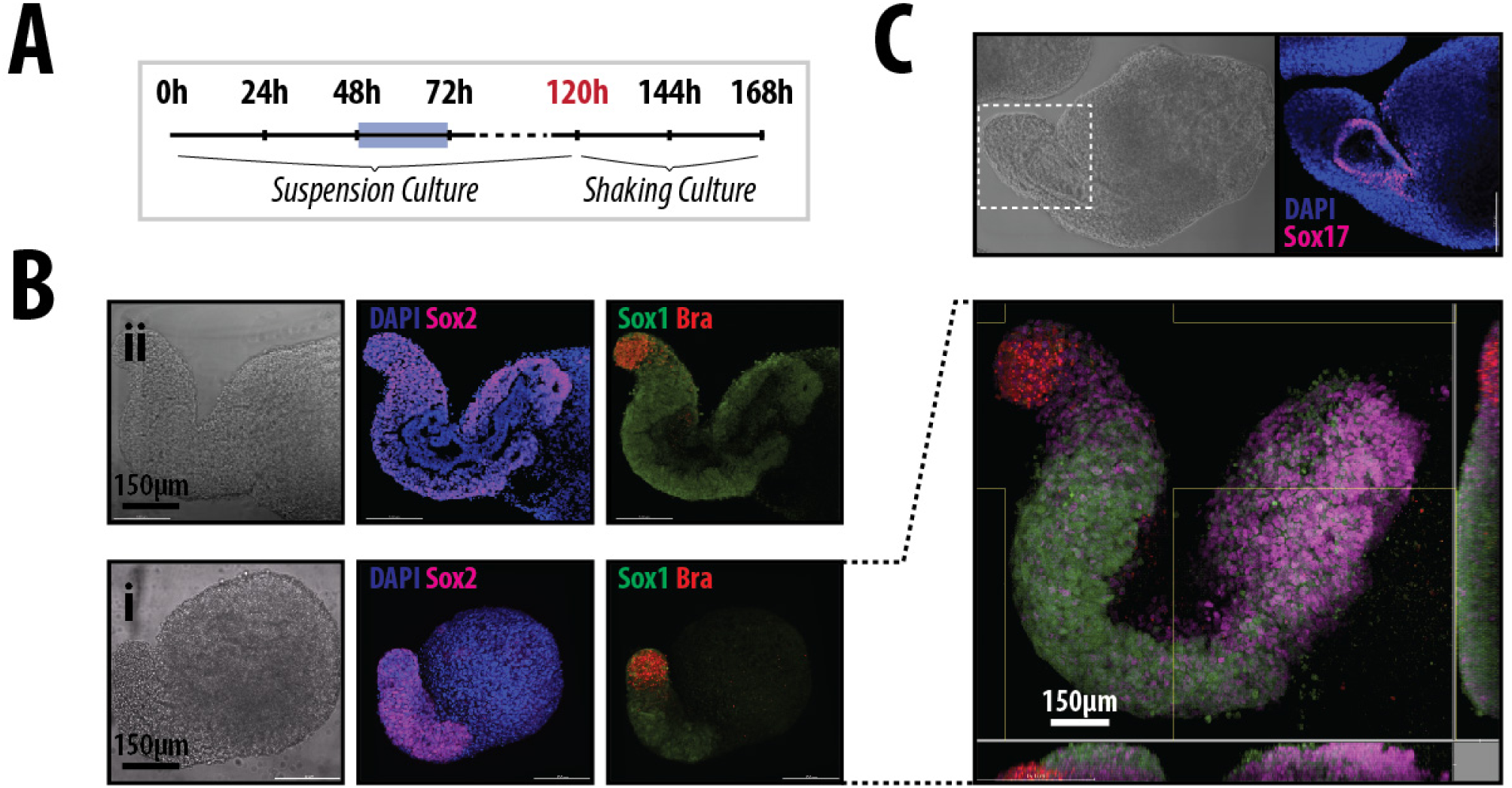
*Gastruloids* in shaking culture develop complex interior structures reminiscent of the primitive gut. (A) *Gastruloids* are maintained in normal suspension culture from 0-120h with a pulse of Chi between 48 and 72h. *Gastruloids* are then shaken in the incubator from 120h-168h (see materials and methods). (B, C) Bright-field and immunofluorescence images of *Gastruloids* fixed at 144h (Bi, C) and 168h (Bii), and stained with DAPI (Blue) and Sox2 (magenta), co-expressing Sox1::GFP (Green) and T/Bra::mCherry (red). 3D reconstruction of the 168h *Gastruloid* reveals the spatial arrangement of Sox1, Sox2 and T/Bra as well as complex internal structures which expressing markers suggestive of a primitive gut tube.

Taken together, these results suggest that by 120h AA, Chiron treated *Gastruloids* have an organisation that resembles the posterior region of the embryo and exhibit its three primary axes. The lack of Sox1 expression at the most anterior region suggests that *Gastruloids* lack brain or head structures and, in this sense, they are very similar to gain-of-function *β*-Catenin mutants (25–27), consistent with their having been exposed to high levels of Wnt signalling in the early stages of development.

### 2.2 Wnt/*β*-catpole of expression is less clearenin signalling provides robustness to the polarisation of T/Bra expression

To understand the emergence of *Gastruloid* axial organisation, we focused on the AP axis and monitored the expression of T/Bra over time from the moment of aggregation using a T/Bra::GFP reporter line (28), as well as the patterns of Wnt and Nodal signalling using the reporters mentioned above. Analysis of *Gastruloids* in N2B27 24h AA revealed weak, heterogeneous expression of T/Bra with a proportion of *Gastruloids* already displaying signs of bias towards one pole (Fig. 3A). The proportion of *Gastruloids* expressing T/Bra at this stage varied between experiments but always reflected the reported pattern (table S1). By 48h AA, the expression levels of T/Bra::GFP had risen uniformly across the population and exhibited a more prominent polarisation (Fig. 3A); however continued culture in N2B27 resulted in variations in both the level of expression and the precision of polarisation (Fig. 3A; DMSO). During this period we also observed expression and localisation of both the Wnt (TLC2) and Nodal::YFP reporters (fig. S2A, B), suggesting that the cells are producing ligands for these pathways; inhibitors of these pathways suppress the expression (not shown). Similar to T/Bra::GFP, TLC2 expression is well defined and polarised. However, the Nodal::YFP reporter is much more heterogeneous by 48h and the pole of expression is less clear (fig. S2B), whereas 24h earlier, its expression is slightly higher and broadly homogeneous (data not shown). Addition of Chi or Wnt3a to the medium between 48 and 72h resulted in enhanced levels of T/Bra::GFP expression by 72h AA compared to the vehicle controls (Fig. 2A) which is maintained in all *Gastruloids* at the posterior tip at higher levels than the control (Fig. 3A).

**Figure 3:**
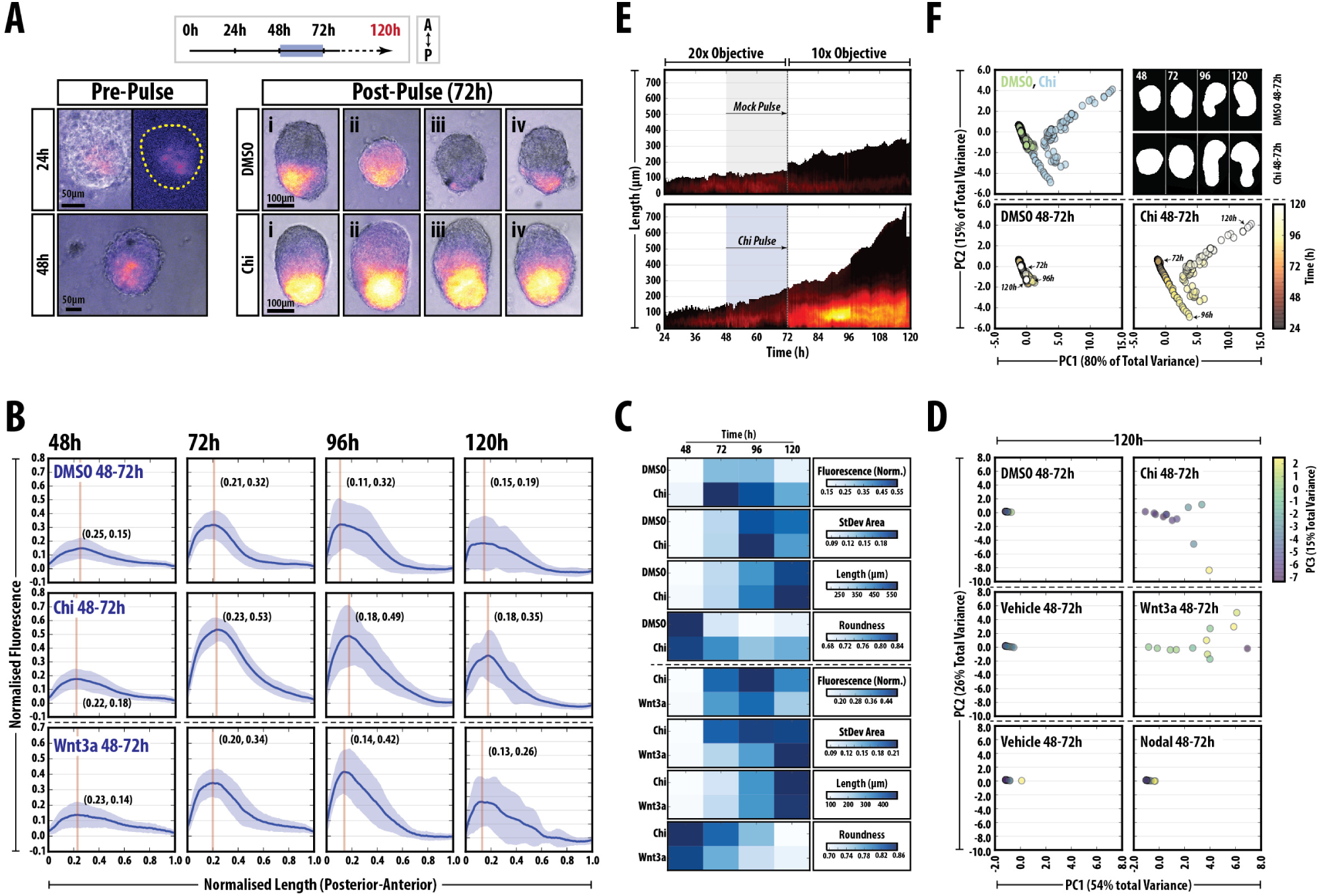
Wnt/*β*-Catenin signalling stabilises and enhances spontaneous symmetry-breaking and polarisation events in *Gastruloids*. (A) Expression of T/Bra::GFP in *Gastruloids* at 24 and 48h prior to the Chi pulse (left), and examples (i-iv) of *Gastruloids* following a DMSO or Chi pulse. Chi stimulation increases the robustness of the response and reproducibility of the phenotype. Experimental design and the AP orientation of the Gastruloids indicated (top right). (B) quantification of T/Bra::GFP reporter expression in individual *Gastruloids* over time from one replicate experiment following DMSO, Chi or Wnt3A treatment as indicated. The maximum length of each *Gastruloid* is rescaled to 1 unit and the fluorescence is normalised to the maximum fluorescence from the Chi condition. Wnt3A condition taken from a different experiment (indicated by hashed horizontal line). Vertical line in each plot marks the peak max and the corresponding coordinates demote the position of this value. (C) Heat maps indicating the average fluorescence (fluorescence norm.), the average area taken up by the standard deviation (StDev Area), average length and the roundness (an indication of *Gastruloid* deformation from spherical) of the *Gastruloids* in the indicated conditions and time-points from the traces in Fig. 3B (Refer to fig. S2 and Materials and Methods for further details). (D) PCA of the *Hu Moments* of the *Gastruloids* in B and C. (E) Live imaging of one representative *Gastruloid* subjected to a pulse of DMSO (top) or Chi (bottom) between 48 and 72h AA. The length of the *Gastruloid* is indicated by the ordinate (posterior =0µm), time on the abscissa and the fluorescence intensity of the reporter in colour. Early time-points (24-72h AA) imaged with a higher power objective. (F) PCA of the *Hu Moments* of a single *Gastruloid* in DMSO or Chi with examples of the shapes at the indicated time-points. Time is indicated by the heat-map.

To garner an understanding of the heterogeneities in the levels of T/Bra::GFP expression over time, we quantified the fluorescence levels of the reporter in a posterior to anterior direction along the spine of the *Gastruloids* (Fig. 3B, C; fig. S2C, D; see Materials and Methods and (18)). We notice that the changes in shape and patterns of gene expression are highly reproducible and this allows us to extract quantitative information about expression and morphogenesis at single-time-points or at regular intervals over time. Exposure of the *Gastruloids* to Chi during 48 and 72h AA results in a tighter distribution of all the measured variables and a higher level of sustained fluorescence than when they are exposed to DMSO (the standard deviation is indicated by the light blue shading about the mean of fluorescence; Fig. 3B, C; fig. S2C). Stimulation with recombinant Wnt3a is able to substitute for Chi and results in less variability in the fluorescence signal and a more rapid acquisition of an elongated morphology (Fig. 3B, C; fig. S2D).

The quantitative analysis allowed us to classify the morphology of the *Gastruloids* in an objective manner through the use of two-dimensional moment invariants, *Hu Moments* (29) in place of the previously acquired shape descriptors. This technique provides the assessment and recognition of two-dimensional geometrical patterns that are independent to the orientation, rotation, size and position (29). The *Hu Moments* of Nodal::YFP *Gastruloids* treated with Chi, Wnt3a or Nodal (with their corresponding vehicle controls) were determined following binary image processing of their bright-field channel and analysed by principal component analysis (PCA; Fig. 3D). Spherical *Gastruloids* from all conditions were found to accumulate tightly, without much variation, and almost all control and Nodal treated *Gastruloids* were found in this region (Fig. 3D). Upon treatment with Chi or Wnt3a, *Gastruloids* developed a degree of elongation which could be well separated with the first principal component (PC1). The second principal component was able to distinguish between Chi and Wnt-treated *Gastruloids*. We also noticed a correlation between PC3 and the elongation potential (Fig. 3D). Using a Sox17:GFP line (30) which reveals endodermal progenitors, we observe that at 120h, Sox17 expressing cells localise anterior to the T/Bra expression domain following the Chi pulse (fig. S3) in the elongating *Gastruloid*, probably creating the progenitors of the tube that can be observed at 160h AA (Fig. 2C).

Live imaging of the T/Bra::GFP reporter throughout the process confirms that Chi enhances an intrinsically polarised expression but also reveals a global transient response to the Chi pulse throughout the *Gastruloid* which relaxes to the original position after the pulse (Fig. 3E; supplemental movie M1,2). The gradual change in morphology of a single *Gastruloid* in either DMSO or Chi conditions can be captured by PCA of the *Hu Moments* (Fig. 3F). The two conditions can be clearly disthave an intrinsic symmetry breaking ability that inguished: DMSO treatment clusters all time-points to one region, whereas over time, Chi treated *Gastruloids* have a well defined trajectory through out the plot (Fig. 3F).

Taken together, these results suggest that *Gastruloids* have an intrinsic symmetry breaking ability that is reflected in the expression of *T / Bra* and made robust and stable by Wnt/*β*-catenin signalling.

### 2.3 Extraembryonic tissues are not required for Axial organisation in Gastruloids

In the embryo, the spatial restriction of T/Bra expression is concomitant with the establishment of the AP axis and the onset of gastrulation at the posterior end of the embryo (31). Genetic analysis has shown that this pattern arises from interactions between signalling systems asymmetrically deployed in the extraembryonic tissues: while the embryo expresses Nodal and, as gastrulation begins, Wnt3 and later Wnt3a, the extraembryonic tissues express and secrete BMP (trophoectoderm) and antagonists for Nodal, Wnt and BMP signalling (visceral endoderm) (7). Interactions between these proteins within the epiblast result in the localisation of T/Bra expression to the proximal-posterior region of the embryo.

To determine the mechanism whereby *Gastruloids* are patterned along the AP axis and compare the process with that taking place in embryos, we first analysed the expression of several genes involved in the AP patterning at 48h AA, when we first observe signs of polarisation in gene expression (Fig. 4A). At this stage, *Gastruloids* expressed *Fgf4, Fgf5, Axin2, Wnt3, Nodal*, and *Cripto*, all of which are expressed in the epiblast in the embryo (Fig. 4A). We also detect low levels of *Lefty1* (Fig. 4A), which in the embryo is expressed mainly in the extraembryonic tissues but also in the epiblast as gastrulation begins. On the other hand, we do not detect expression of genes associated with extraembryonic tissues e.g. *BMP4, Dkk, Furin, Lrp2* and *Dab*, with very low levels of Cerberus (Fig. 4A). By 72h AA in N2B27 we observed increases in expression of *Nodal, Lefty1* and *Fgf5*, decreases in *Fgf4* and the emergence, at low levels, of *Wnt3a* (Fig. 4A). Some of these patterns are Wnt/*β*-Catenin signalling dependent as *exposure to* Chi during 48 to 72h AA leads to a clear increase in *Nodal, Lefty1* and Wnt3a as well as of the Wnt/*β*-catenin targets *Axin2*, *Dkk* and *Cripto*.

**Figure 4:**
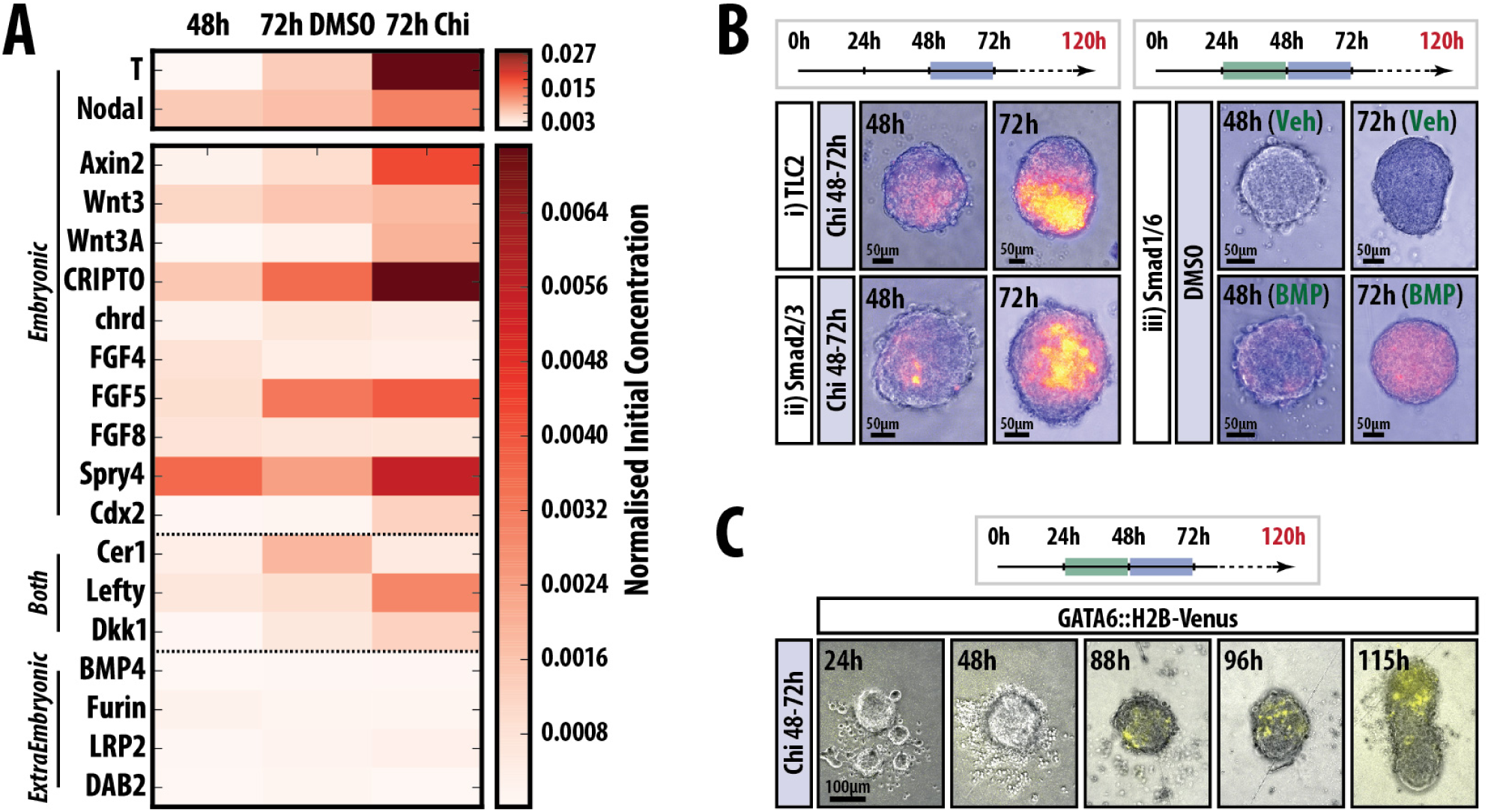
*Gastruloids* do not express genes associated with extraembryonic tissues and progressively activate posterior markers. (A) Quantitative RT-PCR (qPCR) analysis of *Gastruloids* at 24, 48 and 72h AA for genes associated with the Epiblast, Extraembryonic tissues or those expressed in both tissues. (B) Examples of *Gastruloids* expressing reporter genes for the Wnt/*β*-Catenin (TLC2), Nodal (Smad2,3) and BMP (Smad1,5,7) pathways treated as indicated. No BMP signalling is observed under normal conditions (48-72h Chi), but the reporter is responsive to exogenously applied BMP. (C) *Gastruloids* up-regulate the expression of the GATA6::H2B-Venus reporter at ~88h and maintain the expression in the non-elongating (anterior) region over time

These observations support the original contention that *Gastruloids* are made up exclusively of embryonic cells. This conclusion is reinforced by the absence of BMP expression or signalling (Fig. 4B, right), and also by the lack of *GATA6* expression during the first 72h of culture (Fig. 4C), which in the early embryo is associated with the Visceral Endoderm.

The patterns of gene expression at different times AA, together with the timing of the cell behaviours associated with gastrulation that we have described before (14–16, 18), provide landmarks to correlate the development of the *Gastruloids* with that of embryos. They suggest that 48h AA corresponds to the onset of gastrulation in the E6.0 embryo and 72h AA is an approximation to the E7.0.

### 2.4 Nodal signalling promotes T / Bra expression

The expression of signalling reporters suggests that by 48h AA, *Gastruloids* are being patterned through an intrinsic mechanism which relies on Nodal and Wnt signalling (Fig. 3, 4B). To gain insights into this requirement and probe the relationship between the two pathways, we exposed *Gastruloids* to agonists and antagonists of both signalling pathways before or at the time of exposure to Chi. Treatment with the Nodal ALK4 receptor inhibitor SB431542 (SB43) (32) between 48-72h AA in the absence of Chi abolished both the expression of T/Bra::GFP and the elongation, with *Gastruloids* remaining essentially spherical (Fig. 5, fig. S4). Co-treatment with Chi and SB43 (48-72h) severely reduces the levels of fluorescence and greatly impacts the ability of the *Gastruloids* to elongate in a typical manner, with a large degree of variation between experimental replicates (Fig. 5, fig. S4). These results suggest an absolute requirement for Nodal signalling in the expression of T/Bra. To identify a temporal element to this requirement, we pre-treated *Gastruloids* with SB43 between 24 and 48h before pulsing them with Chi (48-72h). These *Gastruloids* are delayed in expressing T/Bra::GFP and the levels show higher degree of variation in terms of the location and expression of T/Bra within individuals; however their ability to elongate is not affected and occasionally enhanced relative to the Chi control (Fig. 5, fig. S4). These results confirm a requirement for Nodal in the expression of T/Bra and suggest that it is possible to separate the axial elongation from T/Bra expression.

**Figure 5:**
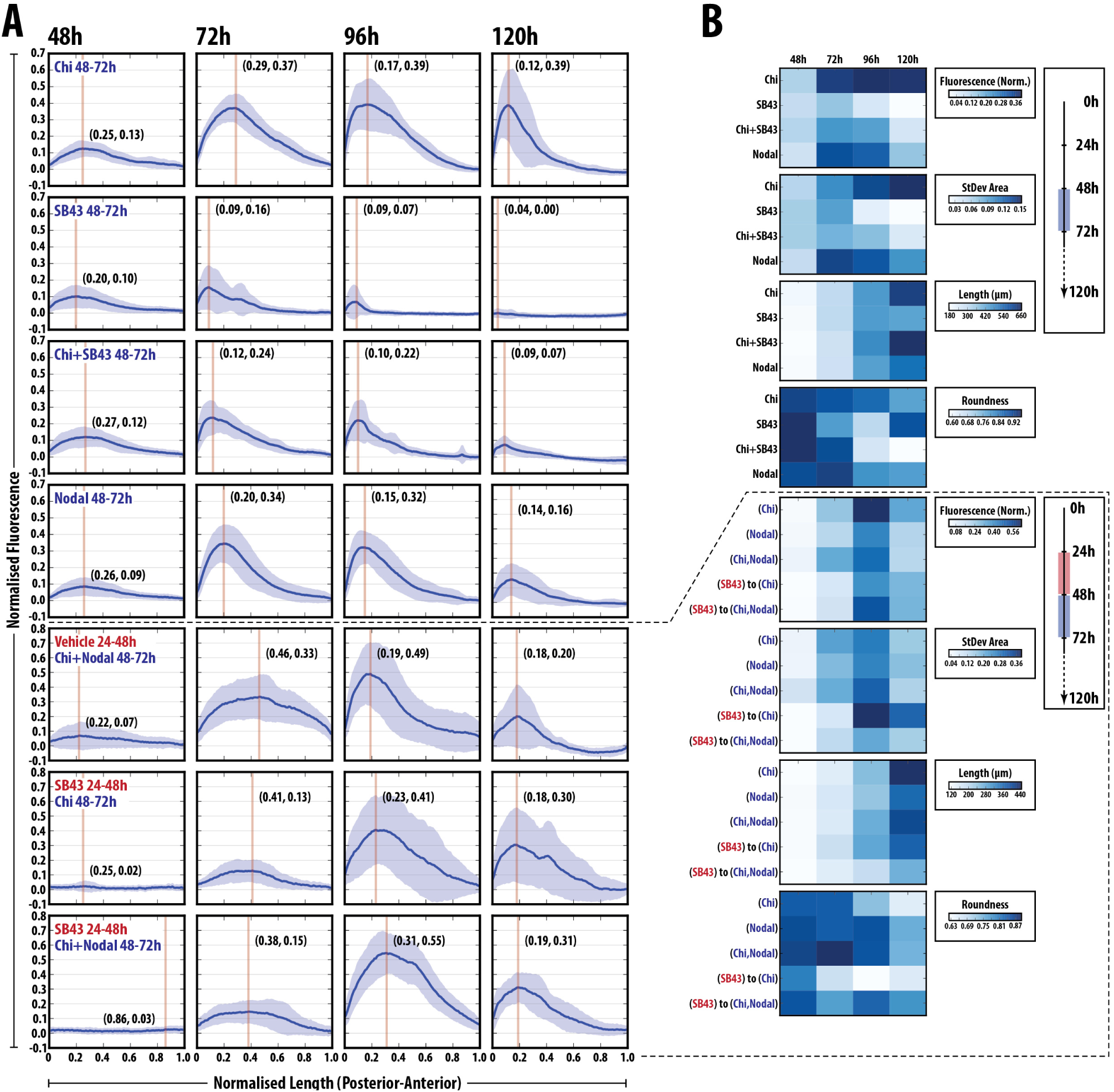
Nodal signalling absolutely required for T/Bra induction and correct patterning. (A) *Gastruloids* stimulated with Chi, SB43, Chi + SB43 or Nodal alone between 48 and 72h AA, or subjected to either vehicle or SB43 pre-treatment (24-48h AA) prior to a Chi, Nodal or Chi+Nodal pulse (48-72h AA). Normalised fluorescence traces shown per condition with the average (blue line) and standard deviation (light blue shading) of *Gastruloid* fluorescence shown following the indicated treatment regimes. (B) The average fluorescence, area of Standard deviation, length and roundness are represented as heat maps (right side). SB43 treatment blocks the expression of Bra::GFP and cannot be rescued by Chi co-stimulation. Inhibition of Nodal signalling has a positive influence on axial length and elongation morphology suggesting that Nodal modulates axial extension (refer to fig. S4 and S5 for further details)

Addition of Nodal alone or with Chiron during 48 and 72h AA results in an increase in T/Bra expression, however the elongation is severely reduced with respect to Chi alone, with *Gastruloids* tending to remain spheroid or ovoid (Fig. 5, fig. S4). This suggests a synergy between the two signalling events. To test this further we tried to rescue the effects of Nodal inhibition on T/Bra expression during 24 and 48hrs. Gastruloids treated with SB43 during 24-48hAA followed by Chi and Nodal co-stimulation between 48 and 72h AA, show enhanced levels of fluorescence compared to Chi and Nodal co-stimulation alone (Fig. 5, fig. S5). Additionally, the increased elongation that was observed with SB43 (24-48h) to Chi (48-72h) treatment is suppressed in this condition, indicating that increased Nodal signalling at this period negatively impacts the elongation, similar to single Nodal stimulation (48-72h; Fig. 5, fig. S5).

These results demonstrate an absolute requirement for Nodal signalling in the expression of T/Bra and its requirement for precise modulation in its levels at specific phases for the elongation. Furthermore, they suggest a negative impact of Nodal signalling on axial elongation.

### 2.5 Wnt signalling promotes T/Bra expression and axial elongation in Gastruloids

To test the role of Wnt signalling in the patterning process, *Gastruloids* were treated in different regimes with either recombinant Wnt3a or its antagonist Dkk1, as well as with small molecule inhibitors of Wnt signalling (IWP3 that affects secretion of all Wnt proteins (33) and XAV939 that increases *β*-Catenin degradation through Tankyrase inhibition (34); Fig. 6). As demonstrated above, Wnt3a is able to substitute more than adequately for Chi during the 48-72h AA period and reduces the fluorescence heterogeneity between individual *Gastruloids* (Figs. 3B, C, A, B). Pre-treatment with Wnt3a prior to a pulse of Chi enhanced the expression of T/Bra::GFP, reduced expression heterogeneity and generated an elongated phenotype more rapidly than controls (Fig. 6). By contrast, pre-treatment with Dkk1, XAV939 or IWP3 before Chi exposure results in a delayed and variable expression of T/Bra (Fig. 6 and fig. S6, S7); however we observe differences in the response to Dkk1 and IWP3, which target Wnt expression and receptor binding, compared to XAV939, that targets active *β*-catenin (Fig. 6, and fig. S6, S7). This suggests a requirement for non-canonical Wnt signalling in T/Bra::GFP maintenance as reductions in Wnt expression (IWP3) or receptor interaction (Dkk1) have a more dramatic effect than reductions in *β*-Catenin activity (XAV939) (Fig. 6). These results reveal that Wnt signalling is essential and the primary signal required for the elongation of *Gastruloids* but that it cooperates with Nodal in the control of T/Bra expression and polarisation.

**Figure 6:**
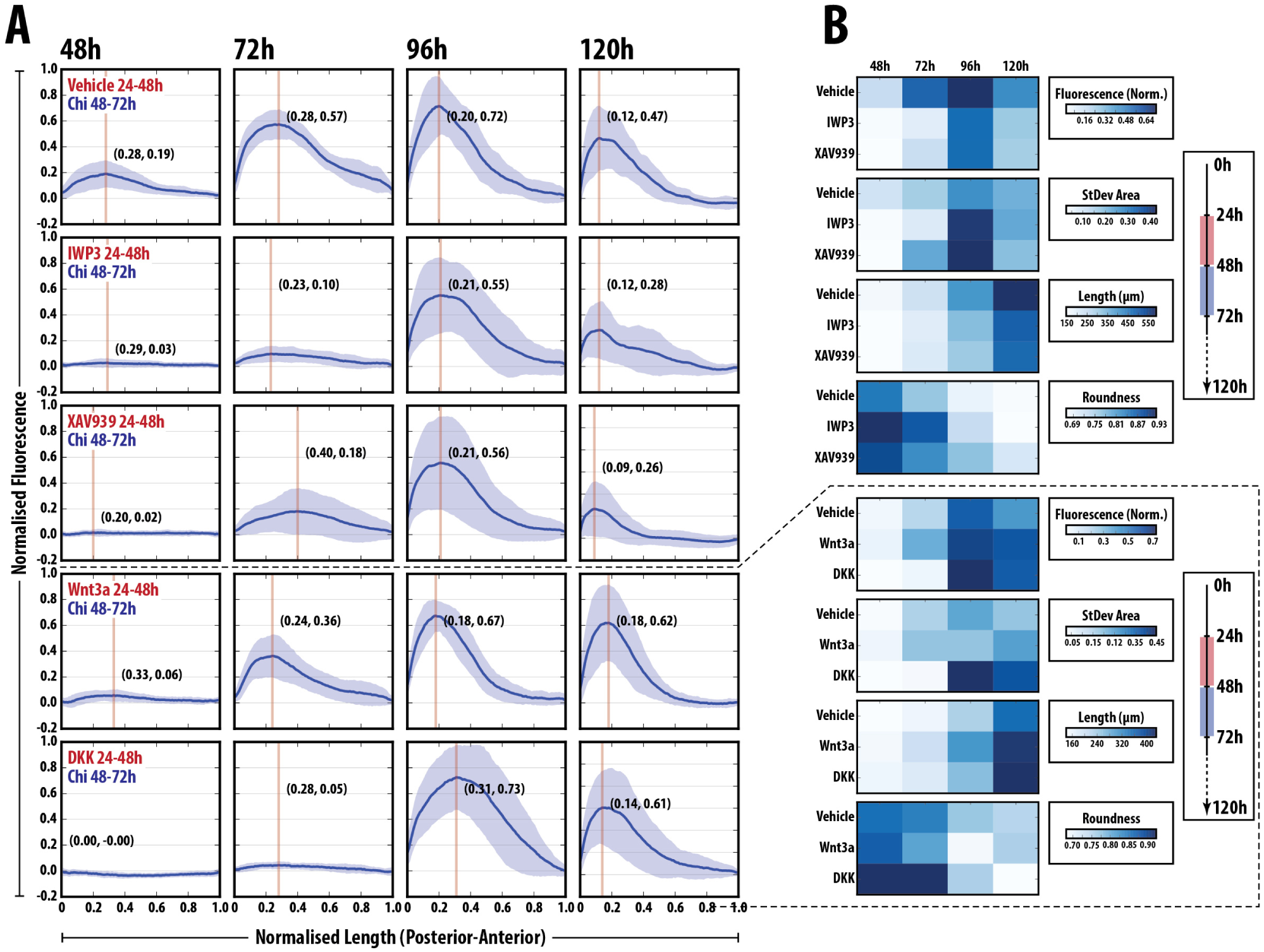
Wnt/*β*-Catenin inhibition delays but does not inhibit Bra::GFP expression. Bra::GRP *Gastruloids* stimulated with a pulse of Chi (48-72h AA) following pre-treatment with vehicle IWP3, XAV939, DKK or Wnt3a. fluorescence traces (A) and heatmaps of the data (B) are shown. Blocking secretion of Wnt proteins with IWP3 effectively abolishes Bra::GFP expression until 96h AA, whereby highly heterogeneous expression is observed. Interestingly, the pulse of Chi can partially rescue Bra::GFP expression at 72h following XAV939 pre-treatment, indicating the requirement for wnt protein secretion in maintenance of expression. Wnt3a pre-treatment reduces the heterogeneity of the response, better defines the pole of expression and maintains high Bra expression for longer than controls (refer to fig. S6 and S7 for further details).

A synergy between Nodal and Wnt signalling during axial organisation has been reported in other organisms (35–37) and is supported by our results which, in addition, suggest different roles for each signalling system. Nodal is essential for the onset of T/Bra expression and Wnt/*β*-catenin signalling does provide amplification and robustness to the response and also promotes Nodal expression in a positive feedback for the process. Our results also demonstrate a requirement for tight temporal regulation of Nodal signalling to allow full elongations to occur.

### 2.6 Wnt/*β*-catenin can generate multiple axes in a Nodal dependent manner

To further delimit the requirements for Wnt/*β*-catenin signalling, we exposed aggregates for 24h at different periods from 24, 42, 48, 52 and 72 h AA and analysed elongation and T/Bra expression (Fig. 7; DAT and AMA: manuscript in preparation). The experiments reveal that the 48-72h period is critical for both the elongation and correct patterning of the Gastruloids. While in all cases there is localised T/Bra::GFP expression and tissue elongation, exposure during the 48-72h period is the most effective (Fig. 7A, B). In the course of these experiments we observed that exposures to Wnt signalling between 24-72 h AA, led to *Gastruloids* with more than one focus of elongation and T/Bra::GFP expression (Fig. 7A, B). In contrast, exposure between 48 and 96 h AA tends to abolish the polarisation of T/Bra::GFP expression and the elongation and lead to *Gastruloids* with overall expression of T/Bra and a rounded shape (Fig. 7A, B). Analysis of the *Hu Moments* (29) of the *Gastruloid* shapes following this treatment regime confirmed that treatment between 48 and 72h AA was most effective in generating an elongated phenotype (Fig. 7A, C).

**Figure 7:**
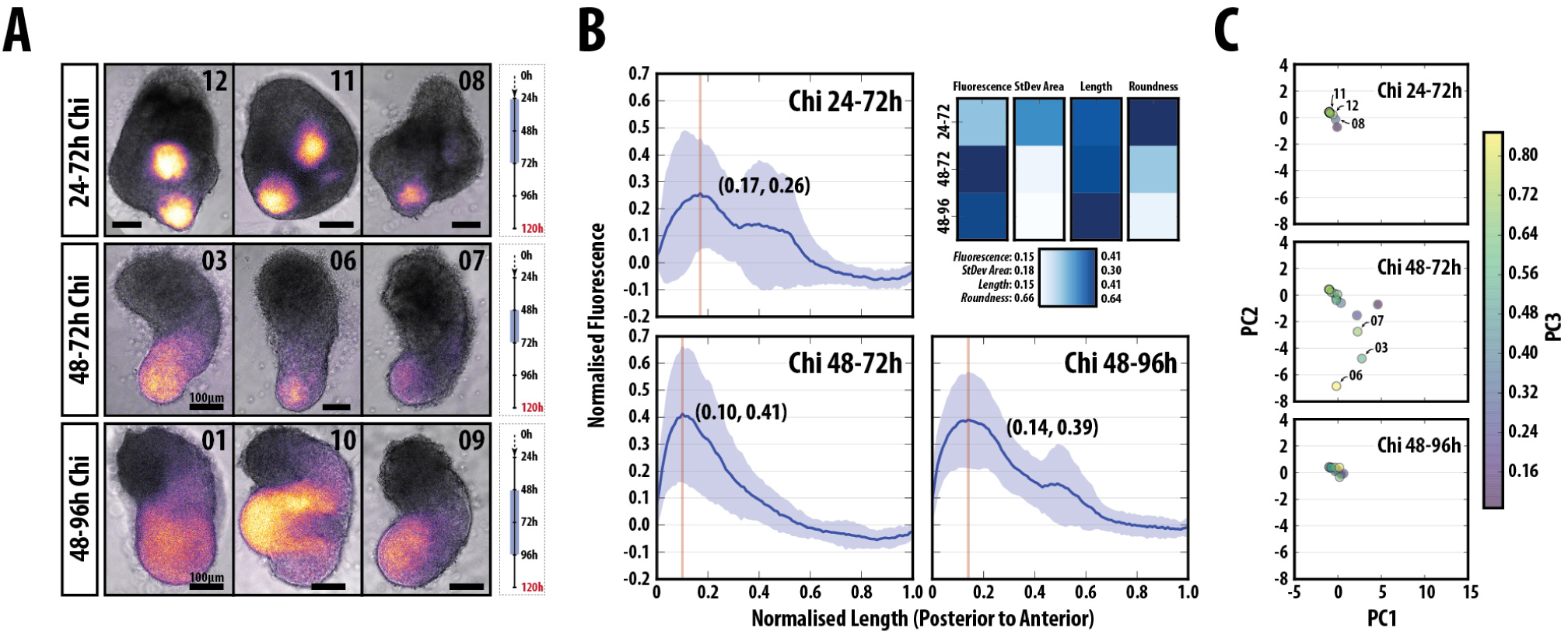
Wnt/*β*-catenin signalling between 48-72h AA is essential for the correct position and expression of T/Bra. (A) Examples of the morphology and expression of T/Bra::GFP *Gastruloids* stimulated with Chi between 24-72h (top), 48-72h (middle) and 48-96h (bottom) AA and (B) the corresponding fluorescence and shape-descriptor quantification. (C) PCA of the *Hu Moments* of the *Gastruloids*. Multiple poles of expression and stunted elongations are observed when Chi is applied between 24-72h AA whereas longer, later stimulation (48-96h) results wider *Gastruloids* and less well defined T/Bra::GFP expression, compared with the 48-72h control.

Together these results reveal two overlapping events centred around the 48h AA time which we have mapped to approximately E6.0 in the embryo. In the 24-48h period there is autonomous axial organisation from within the *Gastruloid* which is stabilised through Wnt/*β*-catenin signalling but critically dependent on Nodal signalling. Following this period (after 48h), it is essential that Nodal signalling is tightly regulated as it negatively impacts the elongation potential of the *Gastruloid* and long exposures abolish elongation without altering the expression localisation of T/Bra::GFP. This highlights the period between 24 and 48h as critical for axial establishment, which is then consolidated in the period after 48h AA.

### 2.7 A polarised source of Nodal signalling is not required for *Gastruloid* patterning

Exposure of *Gastruloids* to Nodal during 48-72h AA does not lead to overall expression of T/Bra suggesting that, like Wnt signalling, a localised source of Nodal may not be required for its effect. We tested this hypothesis using Nodal mutant mESCs (1) (Fig. 8). When aggregated in standard conditions and grown in N2B27 supplemented with the appropriate vehicle controls, Nodal mutant *Gastruloids* remain spherical or ovoid, exhibit a number of protrusions and by 120h AA a large proportion (~90%) have developed small bulbous structures at varying locations (Fig. 8A, B). These *Gastruloids* resemble those treated with SB43 and confirm the absolute requirement for Nodal in symmetry breaking. We then attempted to rescue this *Gastruloids* with various signalling regimes. Addition of Nodal (24-48h AA) reduces the frequency of protrusions but the number is not significantly different from the control (Fig. 8B). Treatment with Chi (48-72h) leads to an increase in the proportion of elongated *Gastruloids* (~50%) supporting a role for Wnt signalling in elongation (Fig. 8A). However the average number of protrusions was similar to controls, with some showing four or more protrusions; the size of the protrusions was also increased relative to the control, but not statistically significant (Fig. 8B). Application of Nodal (24-48h) followed by Chi (48-72h) drastically increased the proportion of *Gastruloids* displaying an elongated, non-protrusion phenotype (0 to 50%) and the number of protrusions was greatly reduced, but not eliminated, compared with the Vehicle to Chi and Vehicle to DMSO controls. Immunofluorescence revealed that Nodal mutant *Gastruloids* treated with Chi were unable to up-regulate the posterior markers T/Bra and Cdx2 compared with previous observations (Fig. 1). However, addition of Nodal prior to the Chi pulse rescued the patterning and location of the reporters (Fig. 8C).

**Figure 8:**
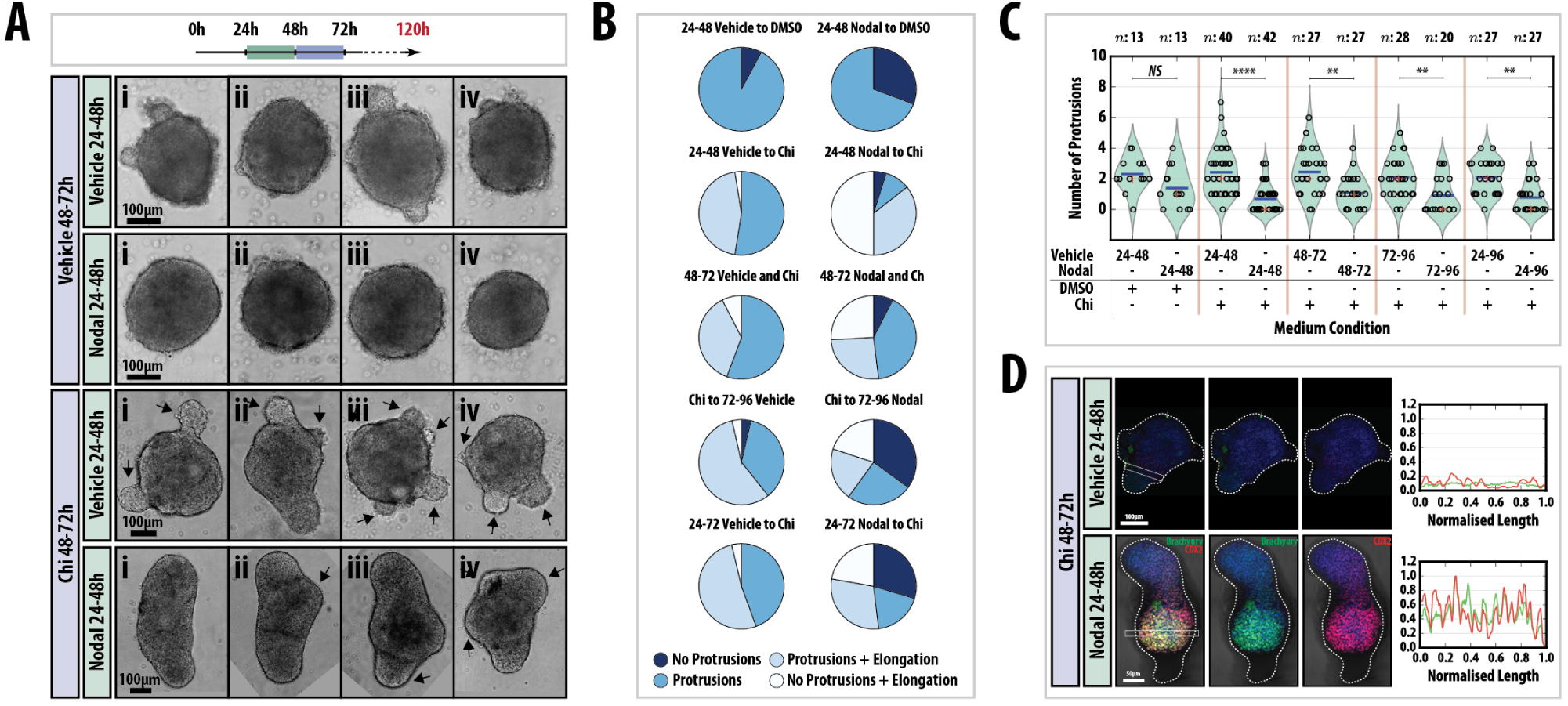
Tight temporal regulation of Nodal signalling is required for axial elongation and proper Axial patterning. (A) Examples of Nodal-/- *Gastruloids* pulsed with either DMSO or Chi (48-72h AA) following a pulse of the vehicle or 100ng/ml recombinant Nodal (24-48h) and (B) the quantification of morphology (pie charts, right). In the absence of Nodal signalling, a number of protrusions are evident which are suppressed by Nodal pre-stimulation, and Chi stimulation enhances an elongated phenotype but does not suppress protrusions. The wild-type phenotype can be rescued if Chi treated *Gastruloids* have been previously exposed to Nodal. Addition of Nodal at different time-points is not able to rescue the elongations (left and fig. S8). (C) quantification of the number of protrusions observed in each experimental condition. Significance determined following Mann-Whitney U test followed by Bonferroni adjustment, comparing selected columns. (D) Immunofluorescence of Nodal-/- *Gastruloids* pulsed with Chi (48-72h) following pre-treatment with vehicle or 100 ng/ml Nodal (24-48h) and stained at 120h with Hoechst (blue), Brachyury (Green) and CDX2 (Red). Nodal addition rescues axial patterning. Later addition of Nodal has less of an effect on the patterning (see fig. S8).

To assess whether the timing and duration of Nodal addition are important for the rescue of the Nodal mutant phenotype, Nodal was applied between 48-72, 72-96 and 24-72h AA, in addition to Chi between 48 and 72h AA and *Gastruloid* morphology assessed at 120h AA (Fig. 8A, right; fig. S8). Although there was some variation between experimental replicates, applying Nodal at later time points reduced the no protrusion-elongated phenotype whilst increasing the No protrusions, no elongation morphology (Fig. 8A, right; fig. S8) compared with the 24-48h Nodal to 48-72h Chi condition. A longer duration of Nodal signalling did not result in effects different from those obtained for 72-96h Nodal.

In summary, a localised region of Nodal expression is not required for *Gastruloid* patterning. However, a tight control of Nodal signalling prior to the Chi pulse is necessary to rescue the Nodal mutants in terms of morphology and AP axis patterning.

## Discussion

We find that *Gastruloids*, mouse embryonic organoids, develop an embryo-like axial organisation in the absence of external patterned influences over a period of five days in culture. significantly they organise themselves in the absence of extraembryonic tissues, which have been shown to drive axial organisation during embryogenesis. This observation leads us to suggest that in vivo, the role of the extraembryonic tissues might not be to induce axial organisation but rather to bias an intrinsically driven symmetry breaking event similar to the one we report here. The deployment of signalling centres around the embryo thus provides a robust source of spatial information that positions the onset of gastrulation in a defined and reproducible location. This ensures that the first cells to exit the primitive streak can easily access the extraembryonic ectoderm, which lies adjacent to them, to form the allantois and primitive blood (38, 39). If the symmetry breaking was stochastic it would be difficult to link gastrulation to the interactions with extraembryonic tissues. Our suggestion is supported by the observation that in the absence of extraembryonic signals, the embryo still exhibits a degree of patterning and axial organisation, though somewhat variable (40, 41).

A most important consequence of the symmetry breaking event in the embryo is the polarised onset of T/Bra expression that will define the initiation of gastrulation and, later, lead to axial elongation (42, 43). In *Gastruloids* this is recapitulated through interactions between Nodal and Wnt signalling that promote the expression and localisation of T/Bra expression between 24 and 48h AA. The activity of signalling reporters in neutral culture medium suggests that ligands for these signalling pathways are expressed in the differentiating cells, something confirmed by RNA expression. Thus the early patterning of the ESC aggregate is driven by intrinsic Wnt and Nodal signalling but its stabilisation requires high levels of Wnt signalling between 48 and 72h AA that boost the expression of Nodal and T/Bra, thus providing a positive feedback for the process. An interpretation of our results is that Nodal provides the initial input on the expression of T/Bra and the organisation of an AP axis but that these effects are enhanced and consolidated by Wnt/*β*-catenin signalling. This possibility is supported by the observation that in the embryo T/Bra expression is initiated and localised in the absence of Wnt signalling, though this pattern is not robust (44). Similar interactions between Nodal and Wnt/*β*-catenin signalling have been described in chick and frog embryos (35–37) and we have also shown them to occur in an adherent culture system of Primitive Streak formation (45). At the molecular level this synergy is supported by reports of molecular interactions between Smad2,3 and *β*-catenin in the regulatory regions of genes expressed in the Primitive Streak and specifically of Nodal and T/Bra (46–48).

Mechanisms to explain how Nodal leads to symmetry breaking during AP axis formation often invoke Reaction-Diffusion mechanisms (49–51). Accordingly, interactions between Nodal and its inhibitor and downstream target Lefty, lead to the asymmetric localisation of both and to the asymmetric expression of target genes e.g. T/Bra. Surprisingly we observe that ubiquitous exposure to Nodal leads to polarisation of T/Bra expression in the aggregate and, moreover, that this will occur when high ubiquitous concentrations of Nodal are provided to a Nodal mutant *Gastruloid*. This observation challenges many of our current notions about patterning driven by Nodal and demonstrates that Nodal needs not be localised to generate an axis. One possible explanation that is consistent with our results is that Nodal signalling initiates the expression of T/Bra but that it is not involved in its refinement and maintenance which depends on a positive feedback between Wnt/*β*-catenin signalling and T/Bra (45). Indeed, several Wnt genes are known to be downstream targets of T/Bra (52) which, in turn, is a target of Wnt/*β*-catenin signalling which thus provide the elements of a positive feedback loop that could be involved in the patterning and localisation of T/Bra and its downstream targets. The absence of a requirement for a localised Nodal signal is supported by the lack of a localisation of Nodal signalling and expression in the *Gastruloids* and also in the embryo, where at the time of gastrulation Nodal is expressed in most of the circumference on the embryo (53).

Our results also highlight that, in addition to, and independently of its role in T/Bra expression and of its interactions with Nodal/Smad2,3 signalling, Wnt/*β*-catenin signalling is central to axial elongation. This provides an independent proof of this well established phylogenetic relationship (54). Timing of exposure suggests two different phases to this involvement. Long exposures to Wnt signalling early (24-72h AA; E5.0-E7.0 in the embryo) can lead to multiple axes, only some of which express T/Bra; this mirrors situations with gain of function of Wnt signalling (55, 56). Increased activity later on (48-96h AA; E6.0-E8.0) results in abolition of the polarity and ubiquitous expression of T/Bra. These observations highlight two temporally separate activities of Wnt. A first one in the establishment and enhancement of the AP axis, probably together with precisely controlled Nodal signalling, followed by a second phase of stabilisation of the pattern and the elongation of the axis. As in the case of Nodal but in a more manifest manner, the observation that a localised source of Wnt/*β*-catenin activity is not necessary for the polarisation of T/Bra expression and the elongation of the *Gastruloid*, questions the widespread notion for a role of Wnt signalling gradients in pattern formation and supports views in which the function of Wnt signalling is to control the signal to noise ratio of events induced by other means (57, 58).

An important feature of the *Gastruloids* is the absence of anterior structures. In this regard they resemble Smad2,3 (59) or Dkk (27) mutants and show that it is possible to orientate an axis without an identifiable head or brain. A likely cause of this deficiency is a combination of the exposure to high levels of Wnt signalling which will suppress anterior development (56, 60) and the lack of a prechordal plate which is essential for neural induction (61). This observation would suggest that while signalling from the extraembryonic tissues might not be strictly necessary for the establishment of an AP axis, it might be essential not only for the reliable positioning of the initiation of gastrulation but also for the location of the brain at the opposite pole.

Over the last few years a number of experimental systems have emerged in which ESCs are spatially patterned and each of them can make a contribution to our understanding of the connection between cell fate assignments and the polarisation of the embryo (62–64). The system that we have developed has some advantages in particular, its reproducibility and robustness allow it to be used in long term studies and screens. Furthermore, the observations that we report here, particularly their ability to be cultured long term and the observation that in addition to the AP and DV axes they can develop an LR asymmetry and generate a node-like structure, suggest that they are a useful model system beyond the early stages of development.

## 4. Supplemental Material

## 5 Materials and Methods

### 5.1 Cell lines and routine cell culture

AR8::mCherry (Smad2,3 reporter) (65), T/Bra::GFP (28), GATA6::H2B-Venus (66), IBRE4-TA-Cerulean (Smad1,5,7) (65), Nodal^condHBE::YFP^ (Nodal::YFP reporter) (24), Nodal^-/-^ (67), Sox1::GFP (23), Sox1/eGFP Bra/mCherry double reporter (SBR) (68), Sox17::GFP (30) and TCF/LEF::mCherry (TLC2) (69, 70) were cultured in GMEM supplemented with LIF, foetal bovine serum, Non-Essential Amino Acids, Glutamax, Sodium Pyruvate and *β*-mercaptoethanol (ESL medium) on gelatinised tissue-culture flasks and passaged every second day as previously described (16, 45, 69, 71, 72). If cells were not being passaged, half the medium in the tissue culture flask was replaced with ESL. All cell lines were routinely tested and confirmed free from mycoplasma.

### 5.2 Immunofluorescence, Microscopy and data analysis

*Gastruloids* were fixed and stained for as required according to the protocol previously described (18). Hoechst3342 was used to mark the nuclei (see table S2 for the antibodies used and their dilutions) except for the images in Fig. 2 where DAPI was used. Confocal z-stacks of *Gastruloids* were generated using an LSM700 (Zeiss) on a Zeiss Axiovert 200 M using a 40 EC Plan-NeoFluar 1.3 NA DIC oil-immersion objective. Hoechst3342, Alexa-488, −568 and −633 were sequentially excited with 405, 488, 555 and 639 nm diode lasers respectively as previously described (45). Data capture was carried out using Zen2010 v6 (Carl Zeiss Microscopy Ltd, Cambridge UK) and data visualisation/analysis performed using Velocity. The z-stacks were acquired of at least 4 *Gastruloids* per condition with a z-interval of 0.5*µ*m for a maximum of 42.2*µ*m. Images were analysed using the ImageJ image processing package FIJI (73).

Widefield, single-time point images of *Gastruloids* were acquired using a Zeiss AxioObserver.Z1 (Carl Zeiss, UK) in a humidified CO_2_ incubator (5% CO_2_, 37°) with a 20x LD Plan-Neflouar 0.4 NA Ph2 objective with the correction collar set to image through plastic. Illumination was provided by an LED white-light system (Laser2000, Kettering, UK) in combination with filter cubes GFP-1828A-ZHE (Semrock, NY, USA), YFP-2427B-ZHE (Semrock, NY, USA) and Filter Set 45 (Carl Zeiss Microscopy Ltd. Cambridge, UK) used for GFP, YFP and RFP respectively, and emitted light recorded using a back-illuminated iXon888 Ultra EMCCD (Andor, UK). Images were analysed using FIJI (73) and plugins therein as previously described (18) and when required, images were stitched using the Pairwise Stitching plugin in FIJI (74). Briefly, the fluorescence intensity was measured by a line of interest (LOI) drawn from the posterior to anterior region of the *Gastruloid* with the LOI width set to half the diameter of a typical *Gastruloid* at 48h (100px with the 20x objective). The background for each position was measured and subtracted from the fluorescence for each *Gastruloid*. Shape-descriptors were generated by converting brightfield images of *Gastruloids* to binary images and measuring them by particle detection in FIJI.

Fluorescence levels were normalised to the maximum obtained in following Chi stimulation, and the maximum length of each *Gastruloid* was rescaled 1 unit. Average fluorescence traces of *Gastruloids* S.D. are shown in the main figures, and the raw data and individual traces in the supplemental data. For live imaging experiments, each well of a 96-well plate containing individual *Gastruloids* were imaged as described above using both the 20x (24-72h) and the 10x (72-96h) objectives, and images captured every 30 min for a maximum of 96h (120h AA). All images were analysed in FIJI (73) using the LOI interpolator (75) with the LOI set as described above.

Data processing, graph plotting, statistical analysis and *Hu Moments* analysis was performed in the Jupyter IPython notebook environment (76, 77) using the following principle modules: Matplotlib (78, 79), NumPy & SciPy (80–82), tifffile (83), Statsmodels (84) and Pandas (85). All code is freely available upon request.

### 5.3 *Gastruloid* culture and application of specific signals

Aggregates of mouse ESCs were generated using an optimised version of the previously described protocol (14, 18). Mouse ESCs harvested from tissue-culture flasks were centrifuged and washed twice in warm PBS. After the final wash, the pellet was resuspended in 3ml warm N2B27 and cell concentration determined using a Moxi™Z automated cell counter with curve-fitting (Orflo Technologies). The number of cells required to generate *Gastruloids* of ~150*µ*m in diameter by 48h (optimised for each cell line, ~300 cells; table S3) was then plated in 40*µ*l droplets of N2B27 in round-bottomed low-adhesion 96-well plates. Counting cells after washing in PBS in this way instead of prior to the washes (as described previously (14, 18)) results in the number of cells required for *Gastruloid* formation being ~100 fewer than previously described as fewer are lost during washing. See table S3 for the number of cells required for each cell line.

In experiments which required the addition of specific factors to *Gastruloids* on the second day of aggregation (24-48h), 20*µ*l medium was carefully removed with a multichannel pipette, and 20*µ*l of N2B27 containing twice the concentration of the required factors was added. This method was preferable to the addition of smaller volumes containing higher concentrations of agonist/antagonists, as the data from these experiments showed more variation between Gastruloids (DAT, PB-J, AMA unpublished). Control experiments showed that replacement of half the medium at this stage did not significantly alter the ability of *Gastruloids* to respond to signals on the third day (DAT, PB-J, AMA unpublished). The next day, 150µl fresh N2B27 was added to each of the wells with a multichannel pipette and left for no more than 30 min to wash the *Gastruloids*; a time delay ensured that sample loss was prevented. Following washing, 150*µ*l N2B27 containing the required factors was then applied. The small molecules used in this study and their concentrations are described in table S4.

To prolong the culture period, individual *Gastruloids* were transferred to low-attachment 24 well plates in 700*µ*l of fresh N2B27 at 120h and cultured on an incubator-compatible shaker for 40h at 50 rpm. 400*µ*l of medium was replenished at 144h and *Gastruloids* were fixed at 160h.

### 5.4 Quantitative RT-PCR

*Gastruloids* (n = ~64 per time-point) from T/Bra::GFP mouse ESCs, subjected to a Chi or DMSO pulse (between 48 and 72h AA), harvested at 48 or 72h AA, trypsinised, pelleted and RNA extracted using the RNeasy Mini kit (Qiagen, 74104) according to the manufacturers instruction as previously described (72). Samples were normalised to the housekeeping gene PPIA. The sequences for the primers are described in table S5.

### 5.5 Orientation of Gastruloids

To define the AP orientation of *Gastruloids*, we have assigned the point of T/Bra::GFP expression as the ‘Posterior’, as the primitive streak, which forms in the posterior of embryo, is the site of T/Bra expression in the embryo (20–22).

### 5.6 Supplemental Figures

**Supplementary Figure 1:**
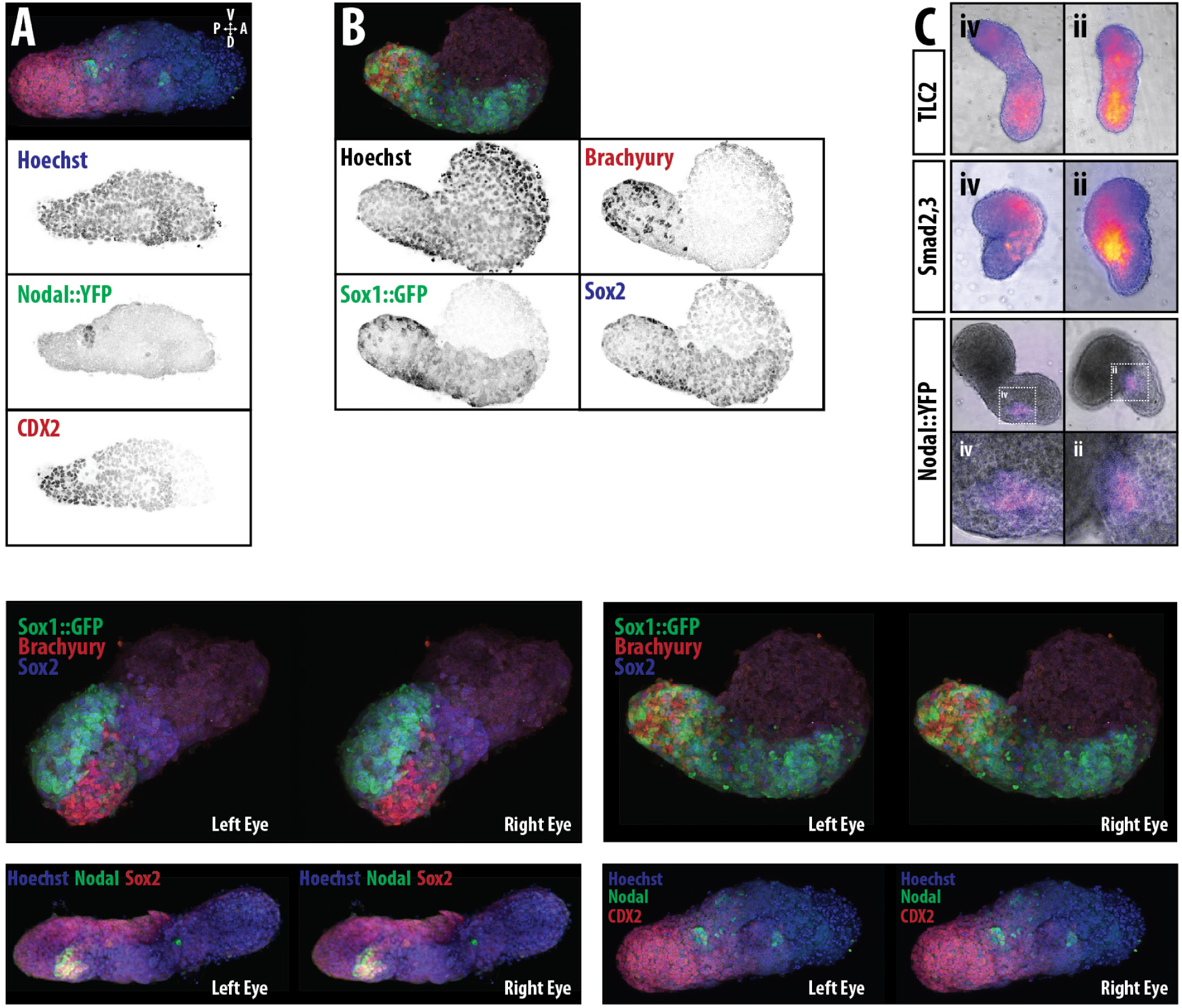
Expression of axial markers in *Gastruloids*. *Gastruloids* from Nodal::YFP (A) and Sox1::GFP (B) mESCs stained for anti YFP (green) and either CDX2 (A) or T/Bra (B) (red) at 120h AA. 3D projection shown above each image. The TLC2 (top), Smad2,3 (middle) and Nodal::YFP (bottom) reporter at 120h (C) are shown as 120h AA. Magnified region for the Nodal reporter indicates a node-like region. (D) Stereo images of the *Gastruloids* from Fig.1A and B.

**Supplementary Figure 2:**
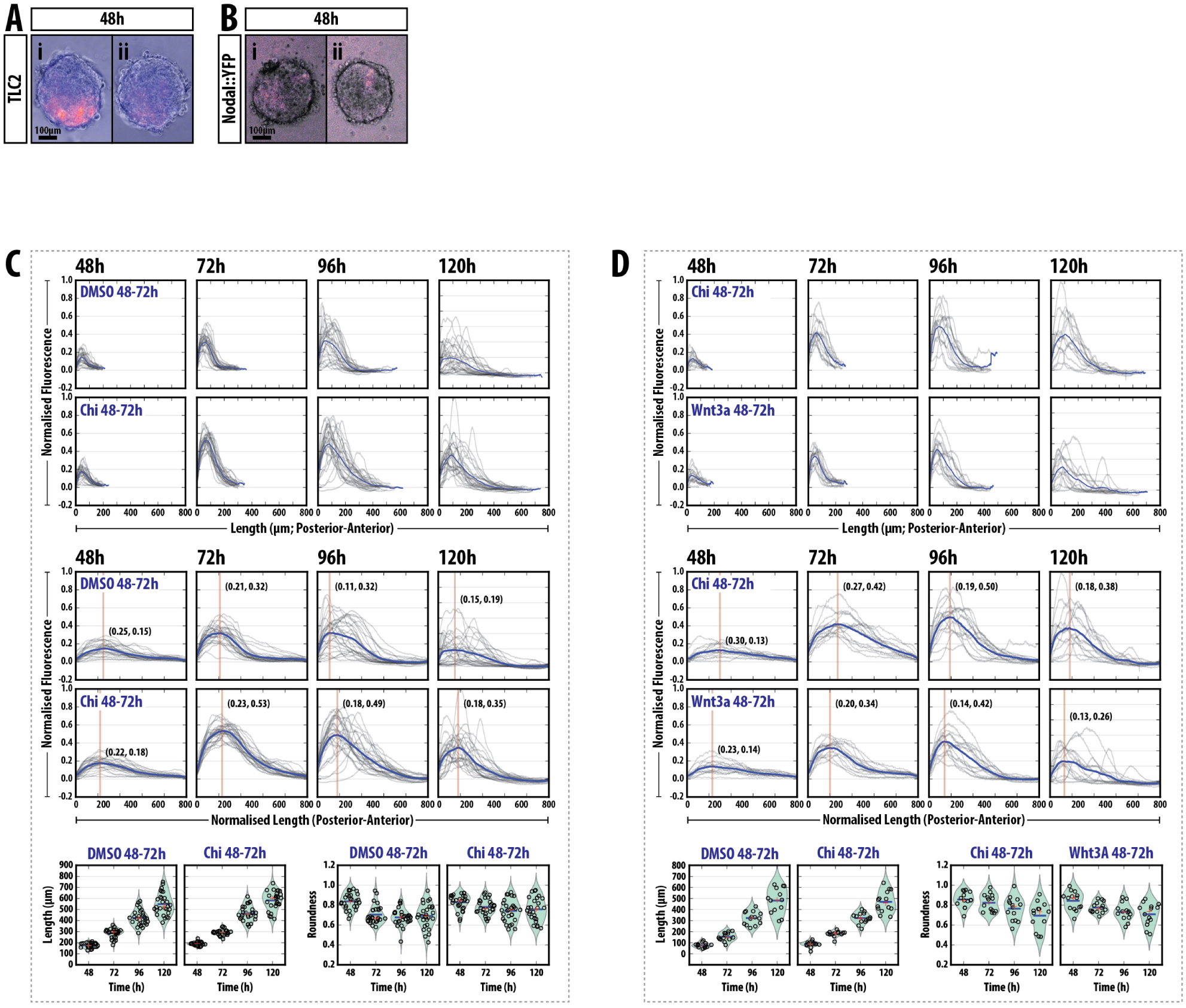
Quantification of *Gastruloid* Fluorescence. Expression of the TLC2 reporter (A) and the Nodal::YFP reporter (B) after 48h in culture. (C, D) Expression of the T/Bra::GFP reporter at the indicated time-points (DMSO or Chi (C) and Chi or Wnt3a (D) stimulation) prior to length normalisation (top) and following normalisation of the length to from 0 to 1 (middle). The bottom panel shows the length and roundness of the *Gastruloids* in the indicated conditions.

**Supplementary Figure 3:**
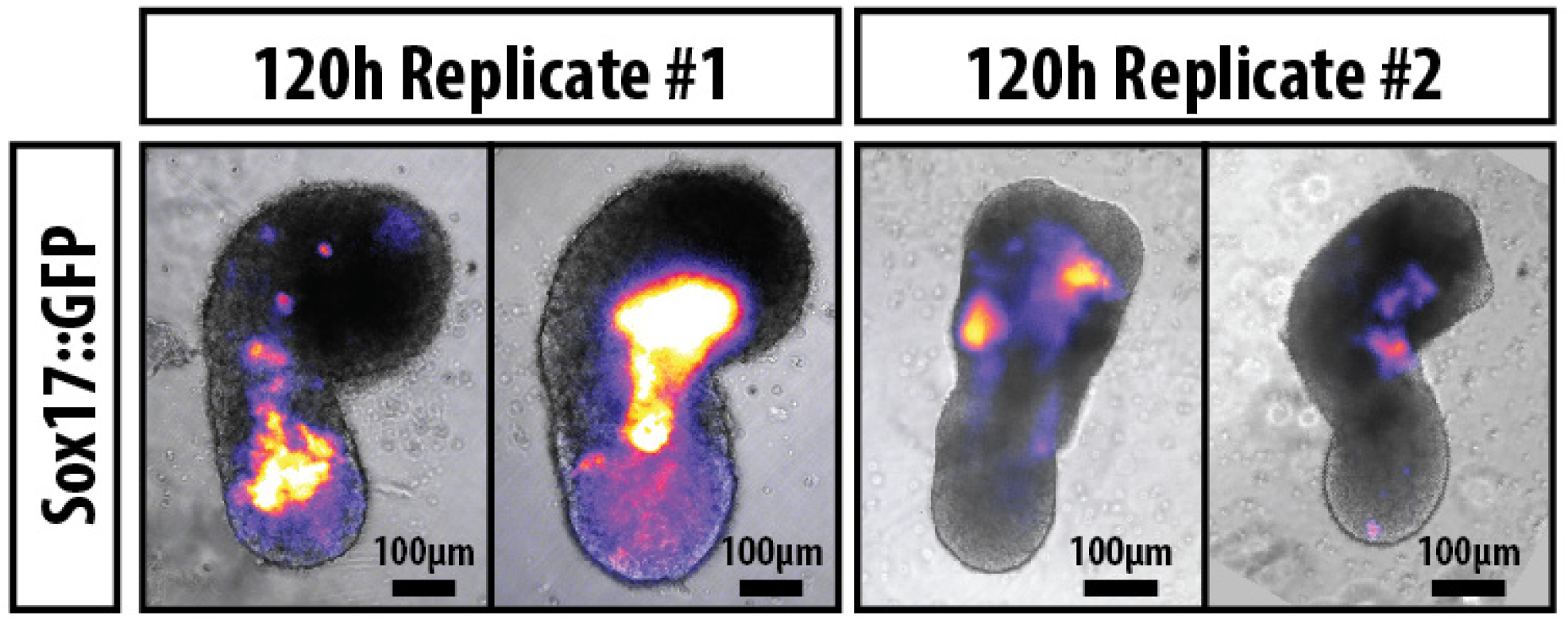
Sox17::GFP is expressed anterior to the elongating region of the *Gastruloids* at 120h AA. *Gastruloids* made from Sox17::GFP mESCs were grown in standard conditions, pulsed with Chi between 48 and 72h AA and imaged by widefield microscopy. Two examples from two replicate experiments are shown, indicating an expression domain more anterior to the elongating posterior region. The posterior of the *Gastruloid* is orientated towards the base of the figure. Scale bar indicates 100μm. Fluorescence levels are not directly quantitatively comparable between experimental replicates due to differences in exposure times and objectives used.

**Supplementary Figure 4:**
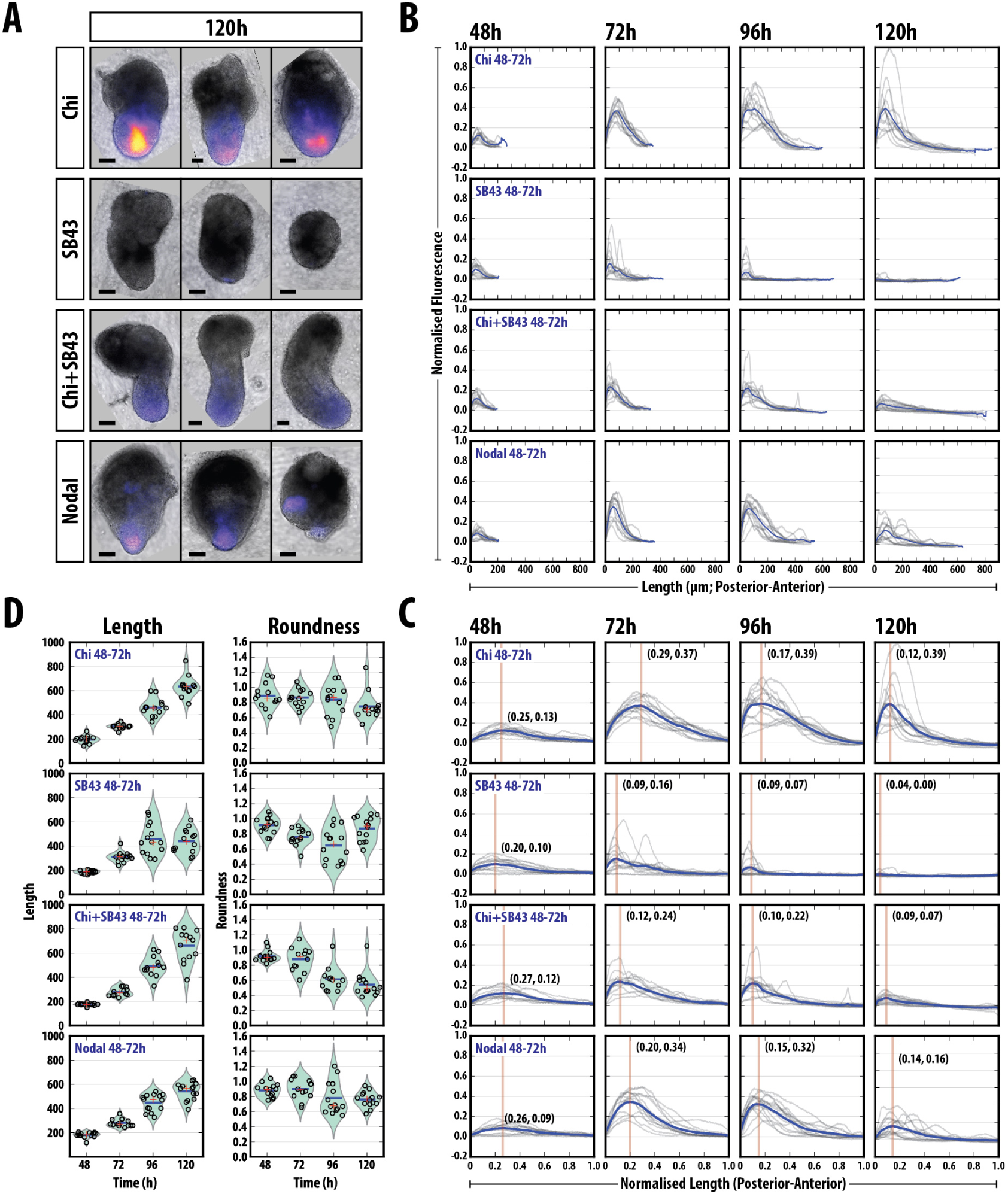
Quantifying the Effect of modulating Nodal signalling in *Gastruloids* (#1). (A) examples of T/Bra::GFP reporter expression in *Gastruloids* treated as indicated. (B, C, D) quantification of the reporter expression at the indicated time-points prior to length normalisation (B) and following normalisation of the length from 0 and 1 (C). The length and roundness of the *Gastruloids* in the indicated conditions (D). Vertical line and coordinates in C correspond to the location and position of the peak maximum. Scale bar indicates 100*µ*m.

**Supplementary Figure 5:**
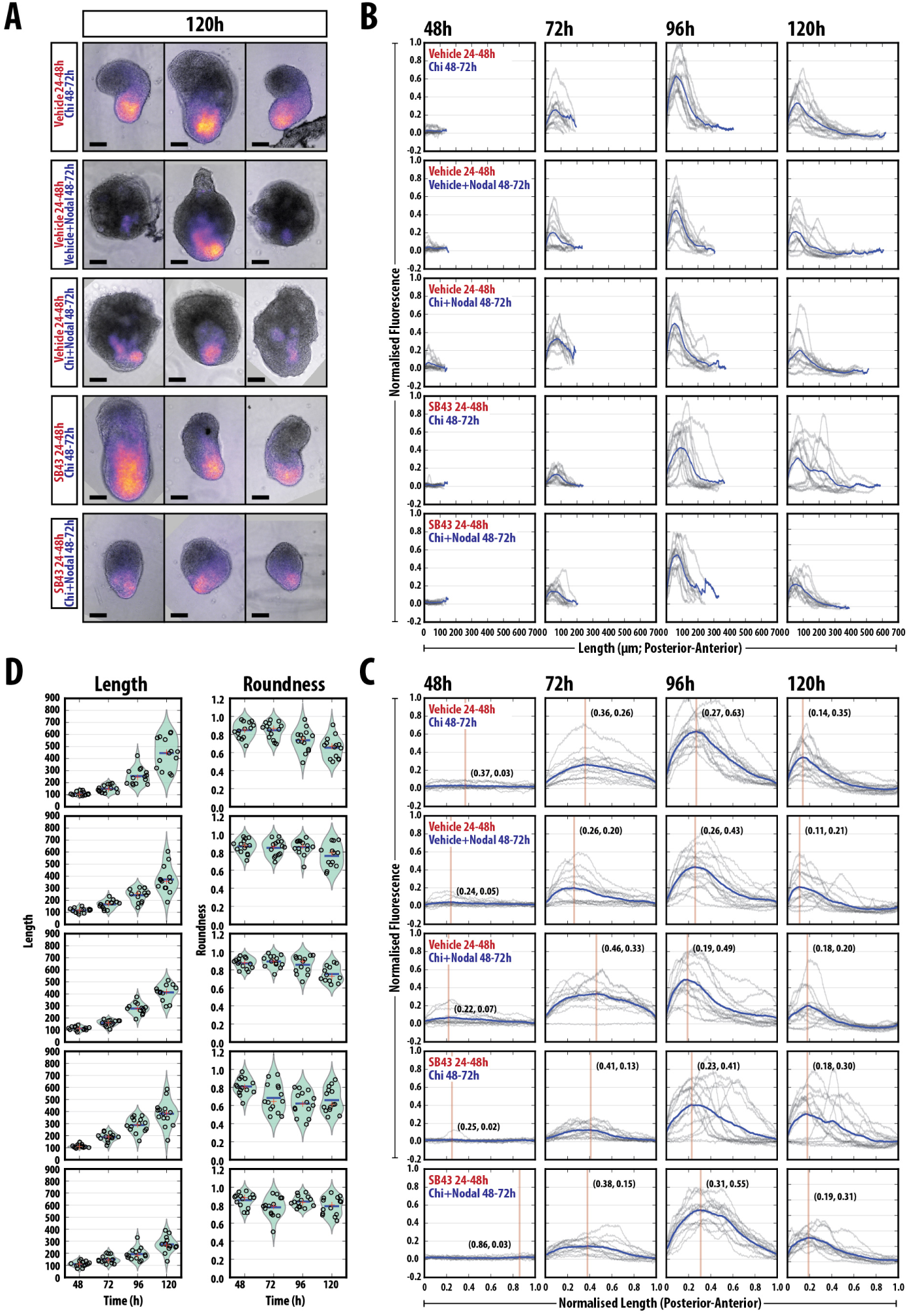
Quantifying the Effect of modulating Nodal signalling in *Gastruloids* (#2). (A) examples of T/Bra: GFP reporter expression in *Gastruloids* treated as indicated. (B, C, D) quantification of the reporter expression prior to length normalisation (B) and following normalisation of the length from 0 and 1 (C). The length and roundness of the *Gastruloids* in the indicated conditions (D). Vertical line and coordinates in C correspond to the location and position of the peak maximum. Scale bar indicates 100*µ*m.

**Supplementary Figure 6:**
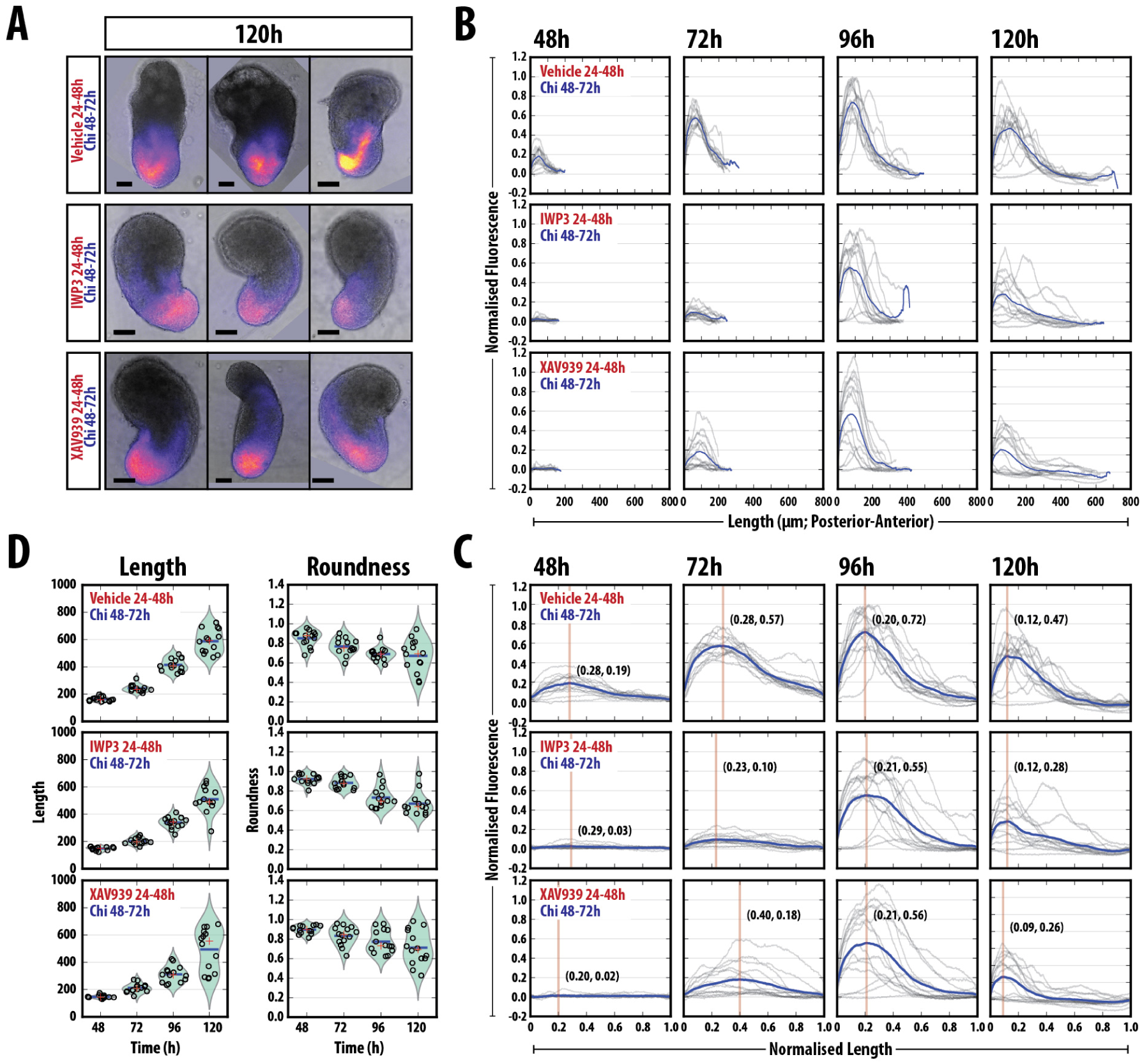
Quantifying the Effect of modulating Wnt/*β*-Catenin signalling in *Gastruloids* (#1). (A)examples of T/Bra::GFP reporter expression in *Gastruloids* treated as indicated. (B, C, D) quantification of the reporter expression prior to length normalisation (B) and following normalisation of the length from 0 and 1 (C). The length and roundness of the *Gastruloids* in the indicated conditions (D). Vertical line and coordinates in C correspond to the location and position of the peak maximum. Scale bar indicates 100*µ*m.

**Supplementary Figure 7:**
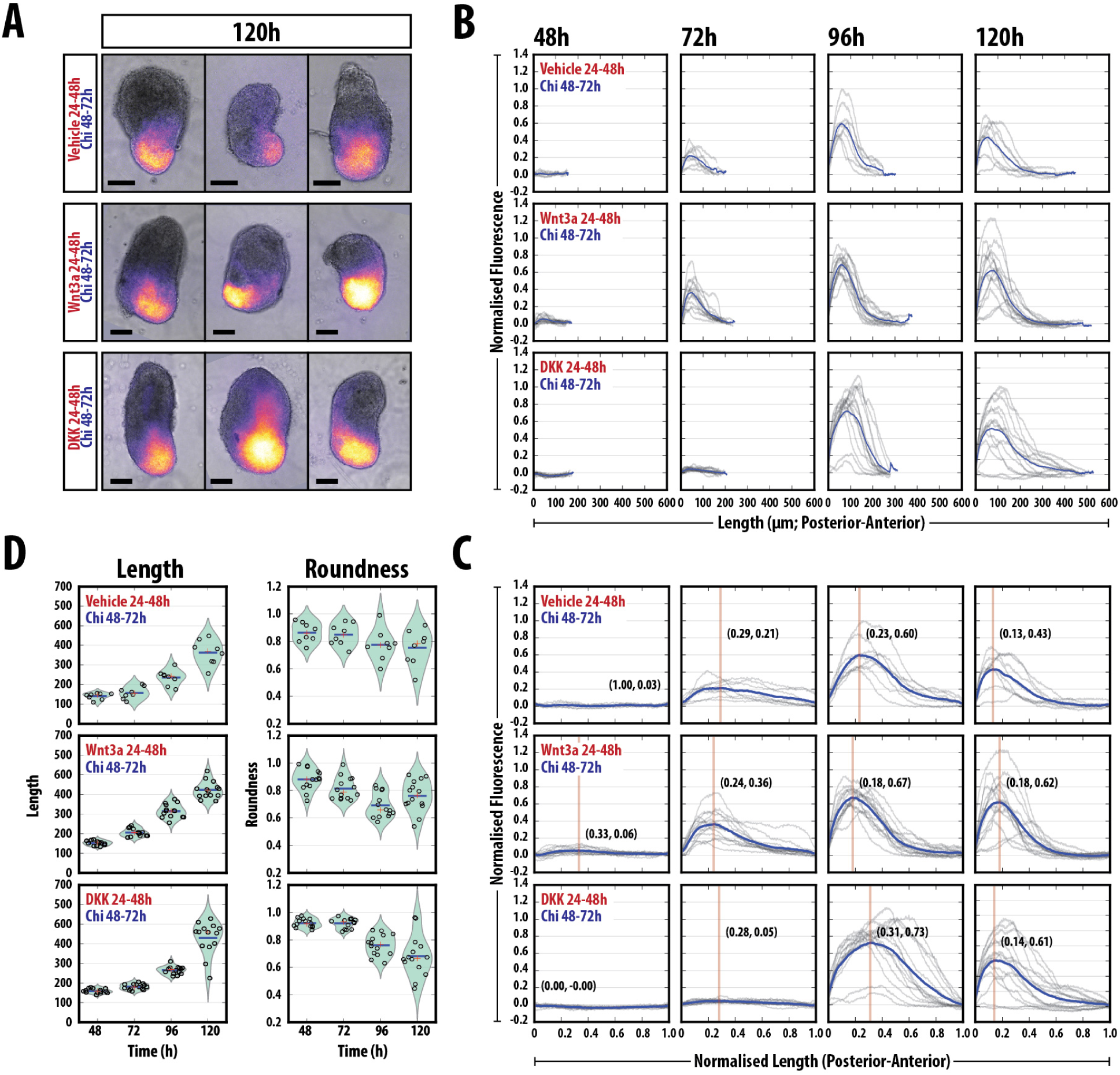
Quantifying the Effect of modulating Wnt/*β*-Catenin signalling in *Gastruloids* (#2). (A) examples of T/Bra::GFP reporter expression in *Gastruloids* treated as indicated. (B, C, D) quantification of the reporter expression prior to length normalisation (B) and following normalisation of the length from 0 and 1 (C). The length and roundness of the *Gastruloids* in the indicated conditions (D). Vertical line and coordinates in C correspond to the location and position of the peak maximum. Scale bar indicates 100*µ*m.

**Supplementary Figure 8:**
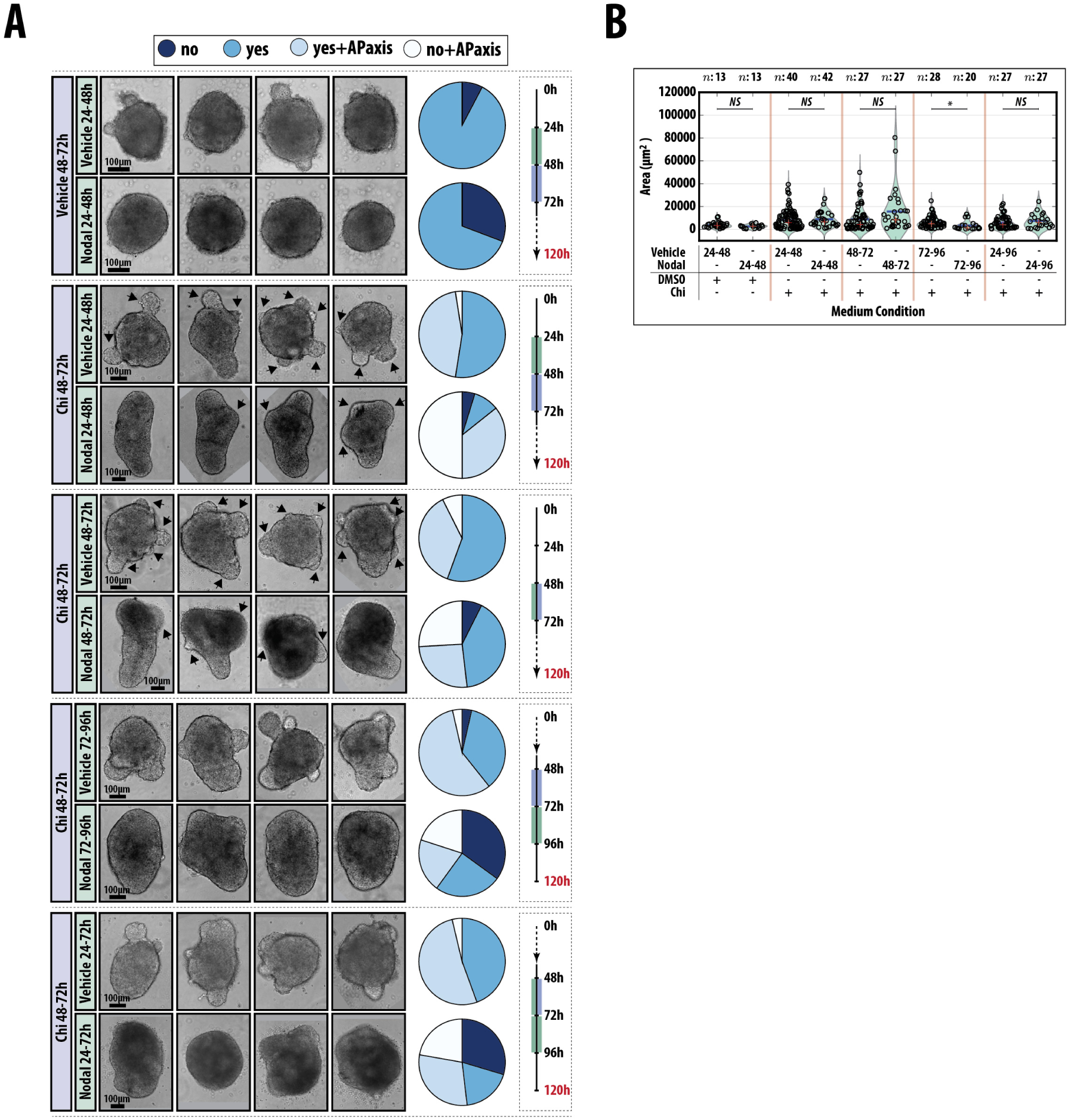
Modulation of Nodal signalling in Nodal mutants. (A) Examples of *Gastruloids* treated with Chi between 48 and 72h with a 24h pulse of either vehicle or Nodal at the indicated time-points (24-48h, 48-72h, 72-96h and 24-72h AA). Pie charts indicated the proportion which do not show protrusions (no), show protrusions (yes), show protrusions with a defined AP axis (yes+APaxis) or dont show protrusions but still have a defined AP axis (no+APaxis). The schematic for the time-course is indicated on the right of the panel. (B) Quantification of the area of the protrusions in the indicated experimental conditions. Significance determined following Mann-Whitney U test followed by Bonferroni adjustment, comparing selected columns.

### 5.7 Supplemental Movies

M1 T/Bra::GFP expression in *Gastruloids* following DMSO treatment (48-72h AA). *Gastruloids* made from T/Bra::GFP mESCs stimulated with a mock pulse of DMSO and imaged by wide-field microscopy from 24h to 120h AA every 20 min. The 20x objective was used between 24 and 72h, followed by the 10x objective from 72h to the end of the experiment. quantification of both the length and fluorescence as a function of time can be seen in Fig. 3E (top).

M2 T/Bra::GFP expression in *Gastruloids* following Chi treatment (48-72h AA). *Gastruloids* made from T/Bra::GFP mESCs stimulated with a pulse of Chi and imaged by wide-field microscopy from 24h to 120h AA every 20 min. The 20x objective was used between 24 and 72h, followed by the 10x objective from 72h to the end of the experiment. quantification of both the length and fluorescence as a function of time can be seen in Fig. 3E (bottom).

### 5.8 Supplemental Tables

**Table 1:**
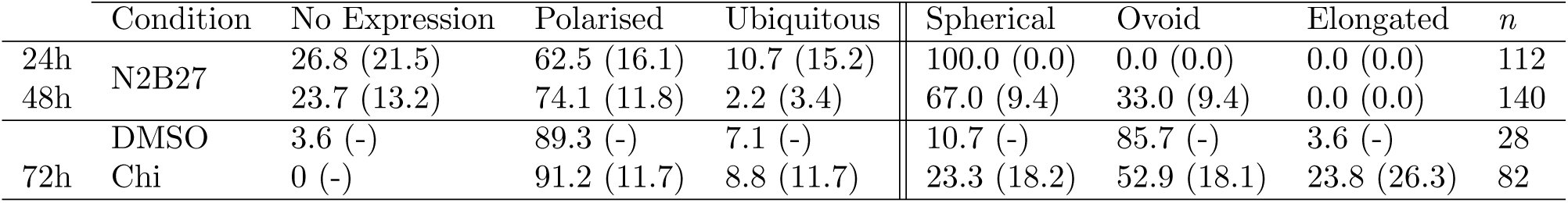
Expression phenotype of T/Bra::GFP mESCs. The proportion of T/Bra::GFP *Gastruloids* not expressing the reporter (No Expression) or displaying either Polarised or Ubiquitous expression at 24, 48 and 72h AA followed by a pulse of DMSO or Chi (72h). The standard deviation is shown in brackets and the number of *Gastruloids* analysed (*n*) are shown.

**Table 2:**
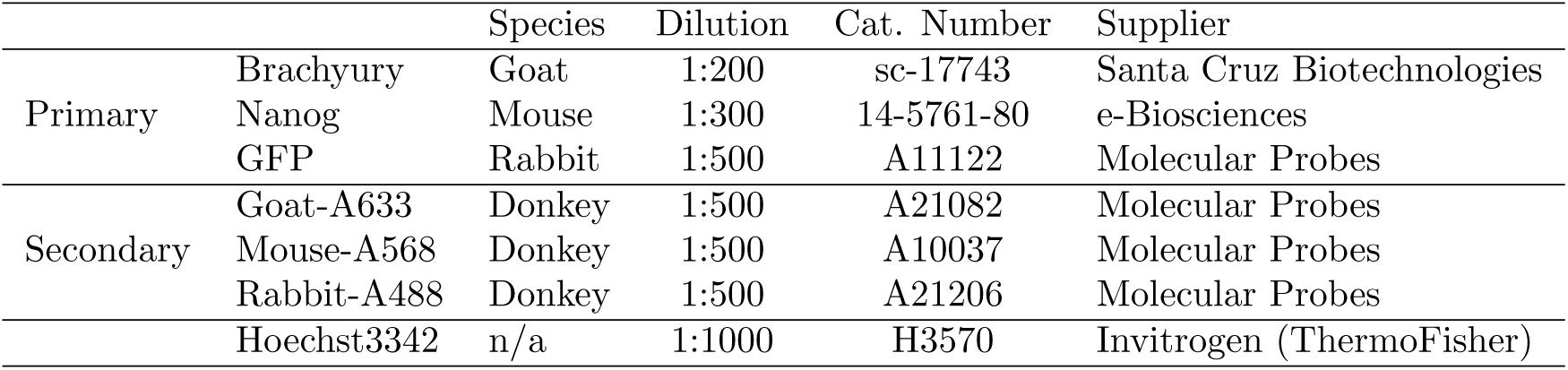
Antibodies and their concentrations used for *Gastruloid* immunofluorescnce with the associated supplier details.

**Table 3:** Cell lines used and numbers of cells required for *Gastruloid* culture. The average diameter of the *Gastruloids* at 48h AA is indicated with the standard deviation and the number of *Gastruloids* measured. N. D.: not done

| Cell Line | Reference | Cells/40μl | 48h Diameter (μm) |
| --- | --- | --- | --- |
| AR8::mCherry | (65) | 450 | 182.7 ± 17.3 (*n* = 83) |
| T/Bra::GFP | (28) | 300 | 161.0 ± 26.2 (*n* = 222) |
| GATA6::H2B-Venus | (66) | 300 | 300 ± (*n* = 10) |
| IBRE4::mCerulean | (65) | 400 | 152.6 ± 12.2 (*n* = 39) |
| Nodal<sup>condHBE::YFP</sup> | (24) | 400 | 138.7 ± 16.1 (*n* = 124) |
| Nodal<sup>-/-</sup> | (67) | 300 | 181.6 ± 23.7 (*n* = 251) |
| Sox1::GFP | (23) | 400 | 112.8 ± 18.4 (*n* = 42) |
| Sox1::GFP/Bra::mCherry | (68) | 300 | N.D. |
| sox17::GFP | (30) | 400 | N.D. |
| TLC2 | (69, 70) | 200-300 | 194.9 ± 20.7 (*n* = 56) |

**Table 4:** Concentrations of Small molecules and recombinant proteins used in this study. Conc: Concentration

| Small Molecule/<br>Recombinant protein | Ref. | Working Conc. | Stock Conc. | Cat. Number | Supplier |
| --- | --- | --- | --- | --- | --- |
| CHI99201 | (19) | 3μM | 10mM | 4423 | Tocris Biosciences |
| IWP3 | (33) | 1μM | 5mM | 04-0034 | Stemgent |
| SB431542 | (32) | 10μM | 100mM | 1614 | Tocris Biosciences |
| XAV939 | (34) | 1μM | 10mM | HY-15147 | MedChem Express |
| BMP4 | | 1ng/ml | 100μg/ml | 314-BP | R&D systems |
| DKK | | 200ng/ml | 100μg/ml | 5897-DK | |
| Nodal | | 1μg/ml | 50μg/ml | 1315-ND-025 | |
| Wnt3a | | 100ng/ml | 40μg/ml | 1324-WN-002 | |

**Table 5:**
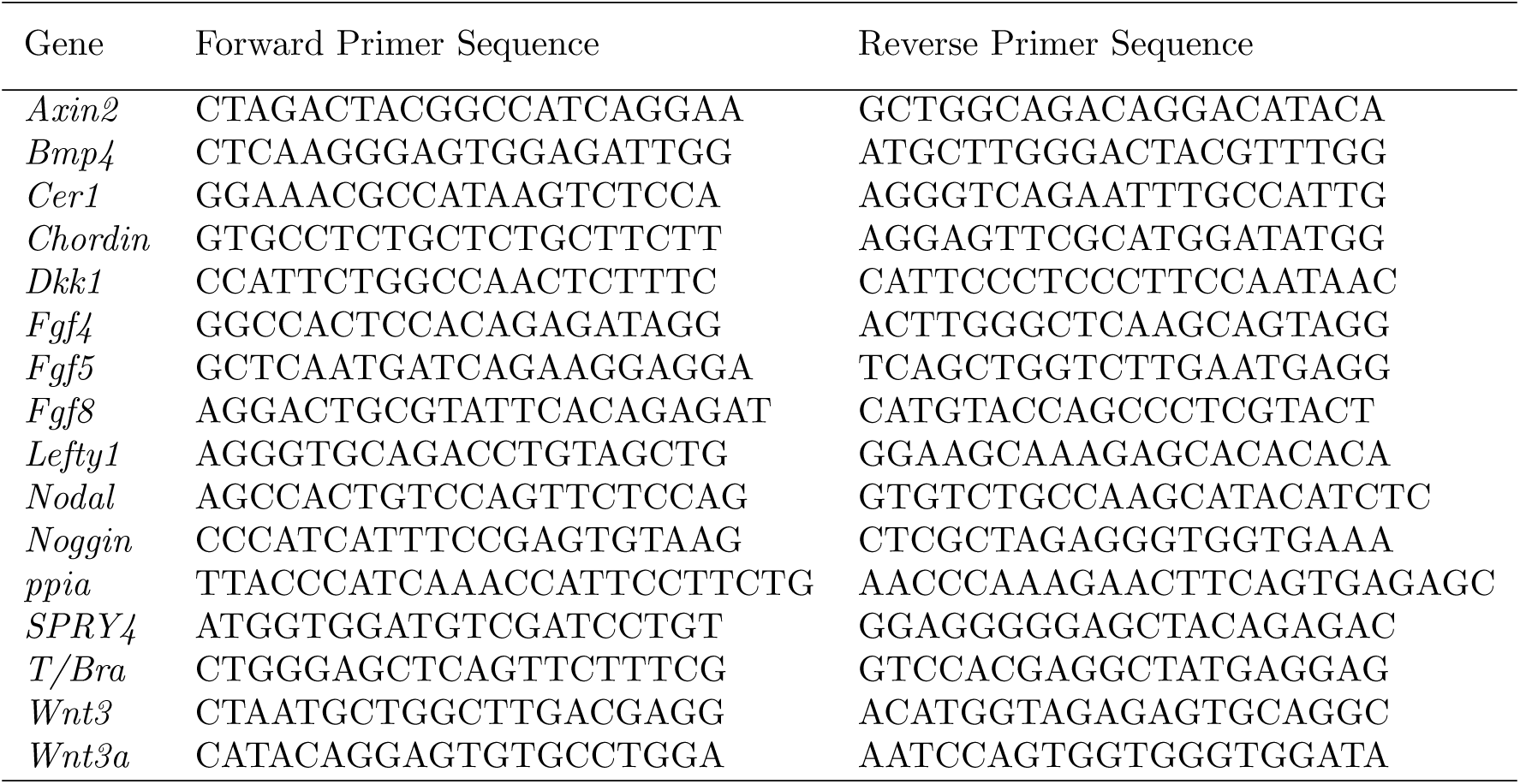
Primer Sequences used for qRT-PCR.

## Acknowledgements

This work is funded by a European Research Council (ERC) Advanced Investigator Award to AMA (DAT, PCH) with the contribution of a Project Grant from the Wellcome Trust to AMA, an Engineering and Physical Sciences Research Council (EPSRC) Studentship to PB-J. Work in the laboratory of ML was funded by support from Ecole Polytechnique Fédérale de Lausanne (EPFL). Portions of this paper first appeared on the BioRxiv pre-print server (15). We want to thank Christian Schröter, Tristan Rodriguez, Kat Hadjantonakis and members of the AMA lab for discussions.

